# The evolution of two distinct strategies of moth flight

**DOI:** 10.1101/2020.10.27.358176

**Authors:** Brett R. Aiello, Usama Bin Sikandar, Hajime Minoguchi, Burhanuddin Bhinderwala, Chris A. Hamilton, Akito Y. Kawahara, Simon Sponberg

**Affiliations:** School of Physics, Georgia Institute of Technology, Atlanta, 30332, GA, USA; School of Biological Sciences, Georgia Institute of Technology, Atlanta, 30332, GA, USA; McGuire Center for Lepidoptera and Biodiversity, Florida Museum of Natural History, University of Florida, Gainesville, 32611, FL, USA; School of Electrical and Computer Engineering, Georgia Institute of Technology, Atlanta, 30332, GA, USA; Department of Electrical Engineering, Information Technology University, Lahore, Pakistan; Department of Biology, University of Florida, Gainesville, 32611, FL, USA; Entomology and Nematology Department, University of Florida, Gainesville, 32608, FL, USA; Department of Entomology, Plant Pathology and Nematology, University of Idaho, Moscow, 83844, ID, USA

## Abstract

Across insects, wing shape and size have undergone dramatic divergence even in closely related sister groups, but we do not yet know morphology changes in tandem with kinematics to support body weight within available aerodynamic power and how the specific force production patterns are linked to changes in behavior. Hawkmoths and wild silkmoths are two such diverse sister families with divergent wing morphology. Using 3d kinematics and quasi-steady aerodynamic modeling, we compare the aerodynamics and the contributions of wing shape, size, and kinematics in 10 moth species. We find that wing movement also diverges between the clades and underlies two distinct strategies for flight. Hawkmoths use wing kinematics, especially high frequencies, to enhance force, but wing morphologies that reduces power. Silkmoths use wing morphology to enhance force, and high amplitude wingstrokes to reduce power. Both strategies converge on similar aerodynamic power and can support similar body mass ranges, but their within-wingstroke force profiles are quite different and linked to the hovering flight of hawkmoths and the bobbing flight of silkmoths. These two groups of moths each fly more like other, distantly related insects than they do each other, demonstrating the convergence and diversity of flapping flight evolution.

## 1. Introduction

The evolution of flapping flight is foundational to the success of many insects [1] and is key for foraging, mate-finding, courtship, predator avoidance, and migration. To conduct these behaviors, significant selective pressure is placed on the flight performance of aerial organisms. Both wing morphology and movement can strongly impact the aerodynamics of a flying animal or machine [2, 3]. Yet, since the origin of flapping flight in insects, a wide diversity of wing morphologies (size, shape, and mechanics), movements, and flight behaviors have evolved [4, 5, 6, 7]. A central question in the evolution of flapping flight is whether divergent strategies (combinations of wing shape, size, and kinematics) arise to achieve similar flight performance. A comparison of disparate species across Insecta shows that not all insects utilize the same flapping flight strategies. Butterflies direct vortices in different directions on upstroke and downstroke [8, 9], while a fly (*Drosophila melanogaster*) and a hawkmoth (*Manudca sexta*) produce lift on both upstroke and downstroke [10, 3, 11]. Mosquitoes use exceptionally high frequency wingstrokes coupled with rapid wing rotations [12] and dragonflies directly actuate the forewing and hindwing independently [13].

Less explored than these disparate comparisons is how closely related sister groups sharing morphologigcal and physiological features diversify to fill different aeroecological niches. Do changes primarily occur in size, shape or kinematics to adapt single flight strategies to multiple behaviors or do all three change in tandem (different overall strategies) when new life histories and behaviors evolve? Considering diversification in closely related, but divergent clades is an opportunity to test how wing size, shape, and movement evolve together and interact to produce species-specific aerodynamics. However, these multiple factors contributing to aerodynamic performance make it challenging to link evolutionary patterns of wing morphology and movement to interspecific differences in flight behavior and aerodynamics.

Hawkmoths (Sphinigdae) and wild silk moths (Saturniidae) are diverse sister clades separated by approximately 60 Ma of evolution [14, 15, 7]. We recently showed wing shape and size diverges between the families through an adaptive shift [7] coincident with the inter-familial divergence in life history [16, 17, 18] and flight behavior [16, 17, 18]. However, wing morphology diverged between the clades in a way that belies simple intuition about how shape should affect performance based on the behavioral demands of each family [7]. Hawkmoths are active, fast-flyers [18], well known for their ability to sustain long duration bouts of hover [19, 20, 21, 22], where some species can track flower oscillations up to frequencies of 14 Hz [22, 23]. Yet, hawkmoths evolved small wings of high aspect ratio (AR), wing loading (*W_s_*), and radius of the second moment of area 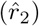, all metrics typically associated with power reduction, efficient force production, and poor maneuverability [7]. In contrast, silkmoth flight behavior is often described as erratic or bobbing [24, 16, 25], which is thought to be advantageous for predator avoidance. Adult silkmoths also lack functional mouth parts and rely entirely on the strictly finite energy stores gathered during the larval period for their entire adult life stage [17]. Yet, silkmoths evolved wing morphology advantageous for maneuverability (low AR, *W_s_*, and 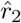), and, despite relying on a fixed energy budget as adults, the clade did not evolve wing morphology advantageous for power reduction [7]. It remains unclear if these shifts in wing morphology led to shifts in flight performance between the clades and how wing movement evolves in tandem and interacts with wing morphology to generate sufficient force to balance body weight while not exceeding available power.

We hypothesize that each clade has evolved a distinct strategy for flight that relies on correspondingly distinct wing sizes, shapes, and movements. If so, silkmoths and hawkmoths with comparable body masses should have similar wingstroke averaged aerodynamic forces, but may differ in their within-wingstroke patterns of force production. Alternatively, the adaptive shift in wing morphology between the two clades [7] has minimal impact on flight performance. A third alternative is that wing movement is conserved across the two clades and the divergence in wing morphology alone plays a role in flight performance leading to the evolution of divergent flight performance between the two clades. While differences between Saturniidae and Sphingidae in wingbeat frequency and observations of erratic versus hovering flight suggest kinematics play a role, morphology and movement have not been integrated in aerodynamic comparisons of the sister clades. To assess the interplay of shape, size, and kinematics, we first quantify three-dimensional wing kinematics during forward flight from live specimens of five species from each sister-family and combine these with our prior analysis of wing shape and size evolution [7]. We then use a blade element model [10, 26, 11] that can be applied across species to estimate the quasi-steady aerodynamic force production and power requirements and separate the contributions of wing size, shape, and kinematics. As the notoriously agile hawkmoths evolved small wings of shapes advantageous for power reduction, we predict that the evolution of wing movement in this group aids in force production and flight control; both can be accomplished through high frequency wing strokes. We predict that the evolution of silkmoth wing movement reduces the power requirements of flight, which can occur through high amplitude wing strokes [27, 28, 29]. Alternatively, silkmoths have not evolved any means for reducing power, which could be an additional factor contributing to the short silkmoth adult lifespan.

## 2. Materials and methods

Details of the mathematical notation used in this study are in Table S1.

### 2.1. Live specimens

Live specimens of five different species from each sister-family (10 total species) were used in this study (Table S2). Species from the hawkmoth (Sphingidae) family include: *Eumorpha achemon, Amphion floridensis, Hyles lineata, Paonias myops*, and *Smerinthus ophthalmica*. Species from the silkmoth (Saturniidae) family include: *Actias luna, Automeris io, Antheraea polyphemus, Hyalophora euryalus*, and *Eacles imperialis*.

### 2.2. Body and wing measurements and morphometries

Body and wing morphologies were digitized for each live specimen using StereoMorph (version 1.6.2) [30] in R (version 3.4.2) following our previous methodology [7]. For all individuals, total body mass (*m*_t_) was measured directly after the individual was flown.

Morphology was analyzed in MATLAB (version R2018b - 9.5.0.944444) following [7]. To generate a combined wing shape from the overlap of the fore- and hindwings, the forewing long axis was rotated to be perpendicular to the body long axis. In hawkmoths, the long axis of the hindwind was also oriented perpendicular to the body long axis. In silkmoths, hindwing orientation was left in its natural position obtained when the wings are splayed open and the moth is at rest. This position was chosen while reviewing videos of silkmoth flight; the long axis of hindwing is always oriented posteriorly and nearly parallel with the body long axis (see supplemental movies). All wing morphology parameters (see supplement) for aerodynamic analysis were calculated from the combined wing following [31].

### 2.3. High speed recordings of moth flight

Flight experiments were conducted in a 100×60.96 working section of an open-circuit Eiffel-type wind tunnel (ELD, Inc, Lake City, MN). See [32] for tunnel details.

Moths were enticed to fly by providing a wind speed of 0.7 ms^−1^. Flight bouts were filmed at 2000 frames s^−1^ for hawkmoths and 1000 frames s^−1^ for silkmoths using three synchronized high-speed cameras (resolution: 1280×1024) (Mini UX 100; Photron, San Diego, CA, USA). The wind tunnel was illuminated with six 850Nm IR light (Larson Electronics, Kemp, TX, USA) and a neutral density filtered white LED “moon” light (Neewer CW-126) [22]. Room lights were also turned on for the diurnal species, (*A. floridensis*).

Videos were digitized and calibrated in XMALab [33]. A total of seven landmarks were digitized on the moth: rostral tip of the head (between the antennae), junction between the thorax and abdomen, caudal tip of the abdomen, left and right forewing hinges, right wing tip, and a point on the trailing edge of the wing (See Table S3 for individual kinematics).

### 2.4. Blade element model summary

A third-order Fourier series was fitted to all kinematics for every wingstroke. Representative wing shape, size, and kinematics for each species were calculated by averaging the wing shapes and time-varying Fourier-fitted kinematics over all wingstrokes of all individual moths.

Species-specific aerodynamic forces were evaluated using a quasi-steady blade element model, estimating the total aerodynamic force as contributions from translational and rotational motion and the force due to added mass [10, 11, 34, 26] and based on hawkmoth (*Manduca sexta*) lift and drag coefficients [11]. A trim search then found equilibrium conditions to balance force and moment over a wingstroke. Trim conditions are given in Table S4.

Total aerodynamic power is the sum of induced, profile and parasite powers [28]. A detailed formulation of the blade element model is in supplement.

## 3. Results

### 3.1. Hawkmoths and silkmoths evolved divergent wing and body morphology

We quantified several morphological features and compared them to our previous work to ensure that the previously identified inter-clade differences in wing and body morphometrics are consistent. Variables include wing area (*S*), the nondimensional radius of second moment of wing area 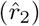, wing aspect ratio (AR), wing loading (*W_s_*), and total body mass (*m*_t_) (Summary data: Table S2).

Hawkmoth and silkmoth wing morphology is well separated in morphospace (Fig. 1A-C). A MANOVA where AR, *W_s_*, and 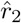 are the response variables and clade is the factor reveals significant differences between clades (F = 34.5 *p*<*0.0001*). AR, *W_s_*, and 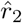 are all greater in hawkmoths than silkmoths by factors of 1.6, 3.7, and 1.1, respectively. Variation in body mass (*m*_t_) spans similar ranges within each family. For a given body mass, *S* is significantly greater in silkmoths than hawkmoths (Fig. 1D) indicated by an ANCOVA between the slopes of each clade (Hawkmoth regression: *r*^2^ = 0.9565, F = 263.7, *p* < 0.0001; Silkmoth regression: *r*^2^ = 0.6273, F = 15.15, *p* = 0.0036; ANCOVA: F = 26.551, *p* < 0.0001). These results recapitulate the broader adaptive shift that we found in wing and body morphology in prior work.

**Figure 1:**
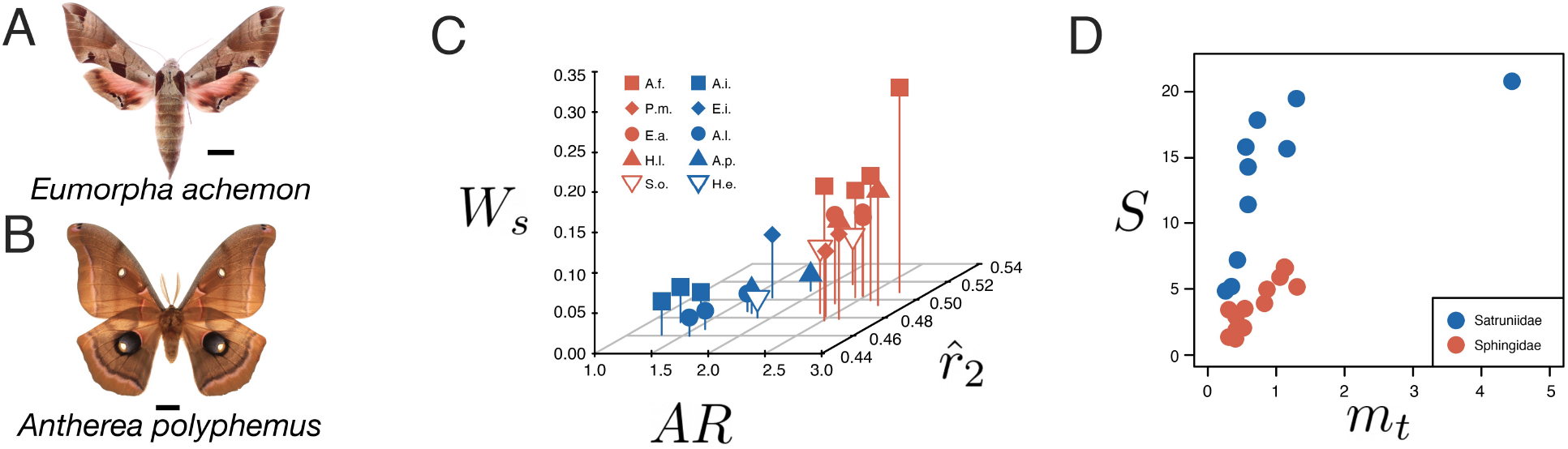
Wing shape and size is divergent between hawkmoth and silkmoth species. (A) A representative hawkmoth (*E. achemon*) and (B) silkmoth (A. *polyphemus*) species used for a detailed comparison of wing kinematics and aerodynamics. (C) Functionally relevant metrics of wing morphology and (D) wing size diverge between hawkmoths and silkmoths.

### 3.2. Hawkmoths and silkmoths evolved divergent wing kinematics

Wing movement can also strongly impact aerodynamics. We next determined if wing movement diverged between the clades. Analysis of 3D kinematics across all ten species reveal strong and consistent inter-clade differences for several variables (Fig. 2; Table S3; recorded kinematics: Hawkmoths, Fig. S4; Silkmoths, Fig. S5). Across the species considered here, hawkmoths fly with a 2.5 times greater average wing beat frequency (*n*) than silkmoths. Silkmoths fly with a 1.2 times greater average sweep amplitude (*ϕ*_p-p_) and 2.3 times greater average inclined (more vertical) stroke plane angle (*β*) than hawkmoths (Table S3). A MANOVA where *n*, 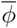, and *β* are the response variables and clade is the factor reveals significant differences between clades (F = 32.546, *p* < 0.0001).

**Figure 2:**
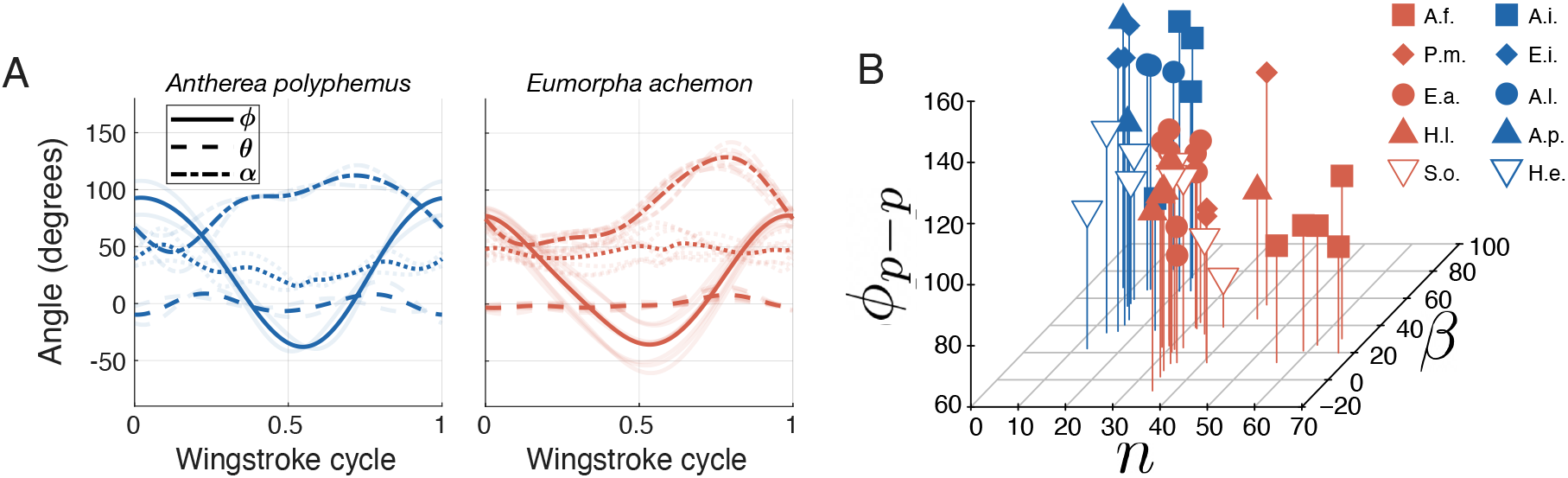
Wing kinematics are divergent between hawkmoths and silkmoths. (A) Within wingstroke kinematics of the sweep (*ϕ*), deviation (*θ*), and feathering angles (*α*) for each of the representative species: *Eumorpha achemon* (Hawkmoth) and *Antherea polyphemus* (Silkmoth). (B) Summary kinematics of wing stroke amplitude (*ϕ*_p-p_), frequency (*n*), and stroke plane angle (*β*) for all 10 species used in this study. The key for species abbreviations for each symbol can be found in Table S1.

Within wingstroke time-varying deviation angle (*θ*) and feathering angle (*α*) differed slightly between hawkmoths and silkmoths (Fig. 2 A-B, S4, S5). On average, silkmoths had 1.5 times larger *θ* amplitude. Whereas, hawkmoths had 1.2 times larger *α* amplitude and 1.1 times smaller mean *α*. These differences are also evident when comparing an exemplar from each clade: *E. achemon* (hawkmoth) and *A. polyphemus* (silkmoth) (Fig. 2 A).

### 3.3. Hawkmoths and silkmoths evolved convergent wing-stroke averaged aerodynamics and power requirements

Given that hawkmoths and silkmoths evolved differences in wing shape, size, and movement, we assessed the overall aerodynamic force and power implications between the two clades. Again, while we focus on two exemplar species (Fig. 3 A), the hawkmoth, *E. achemon* and the silkmoth, *A. polyphemus*, these trends are consistent across the species of each respective family (Fig. S6). Despite the large morphological differences, hawkmoths and silkmoths have overlapping total body masses suggesting that they produce comparable aerodynamic forces at steady state. We found that all species were able to trim to equilibrium conditions using the observed wing stroke kinematic variation in each species and prior observed variation in *C*_L_ and *C*_D_ (Table S4). *Actias luna* was an exception in that its forces trimmed, but a small pitching moment persists in the model. Thus, the combinations of wing shape, size and kinematics, despite being significantly divergent, produce comparable wingstroke-averaged aerodynamic forces (Fig. S6).

**Figure 3:**
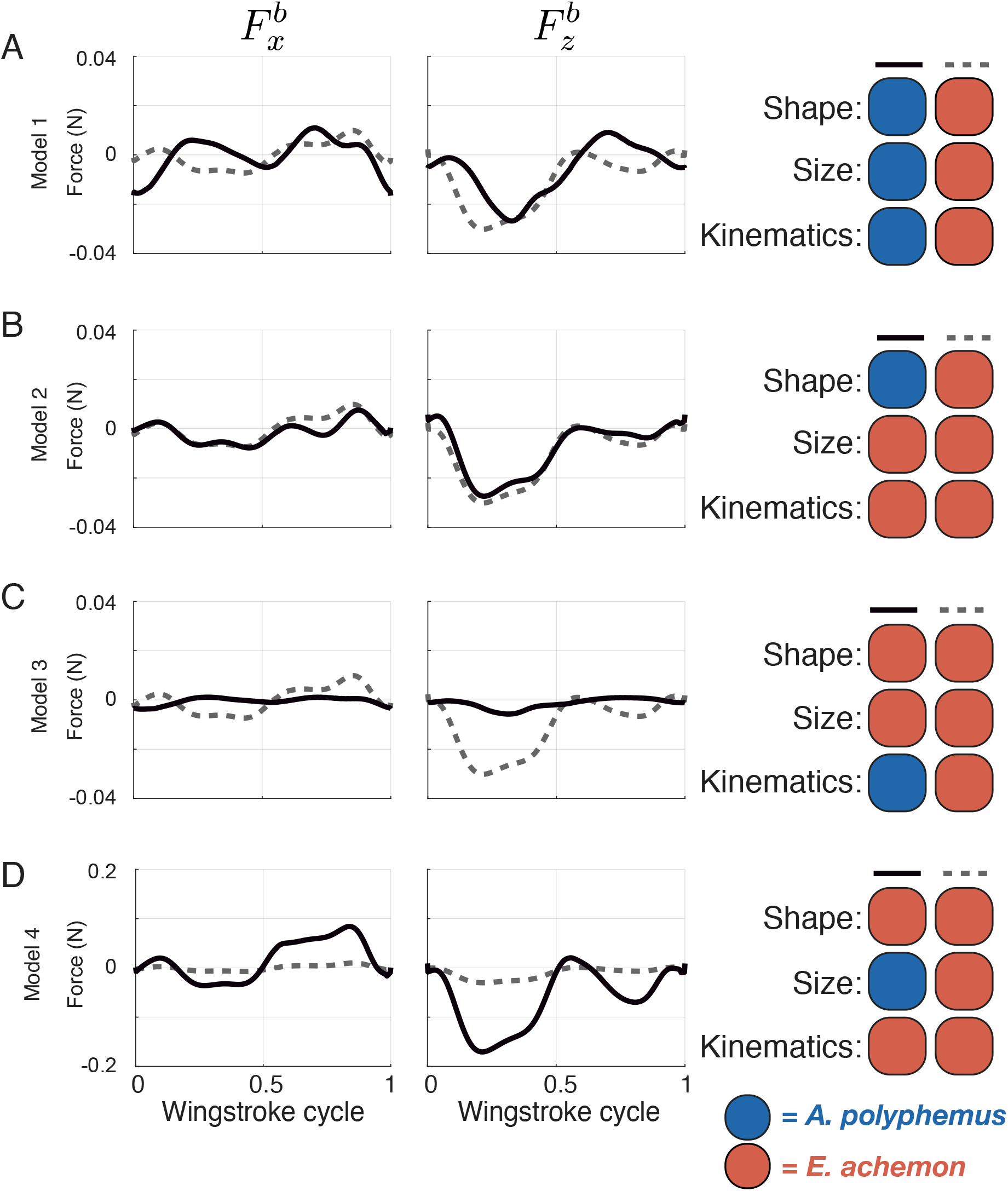
Quasi-steady aerodynamic force production by the right wing in the body-centered coordinate system. The two wings of each model are identified by solid and dashed lines, respectively. The dashed line is the same in each model. The colors on the right side indicate the specific morphology and movement parameters used for each model. Red represents variables from *A. luna* and blue represents variables from *E. achemon*. (A) Model 1 compares interspecific aerodynamics between A. *luna* (solid line) and *E. achemon* (dashed line). Models 2 (B), 3 (C), and 4 (D) investigate aerodynamic impact by changes in wing shape, movement, and size, respectively. All forces are only presented for a single right wing. The negative 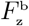 direction points upward and the positive 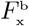 direction points forward.

Similar average aerodynamic forces do not imply similar aerodynamic power across clades. Nonetheless, the power requirements of forward flight are also consistent within and between each clade. Total body mass-specific aerodynamic power requirements range from 15.06 to 18.32 W kg^−1^ in hawkmoths and 11.76 to 17.18 W kg^−1^ in silkmoths (Fig. 4 A). However, there were differences in which component of aerodynamic power contributed greatest. Induced power (*P*_ind_) was always the largest contributor to total power in hawkmoths and 3 silkmoths species, but in two silkmoths (*A. luna* and *A. polyphemus*) profile power (*P*_pro_) was greater than *P*_ind_ (Fig. 4 A). Parasitic power (*P*_par_) was negligible across all species (Fig. 4 A), consistent with prior studies [28].

**Figure 4:**
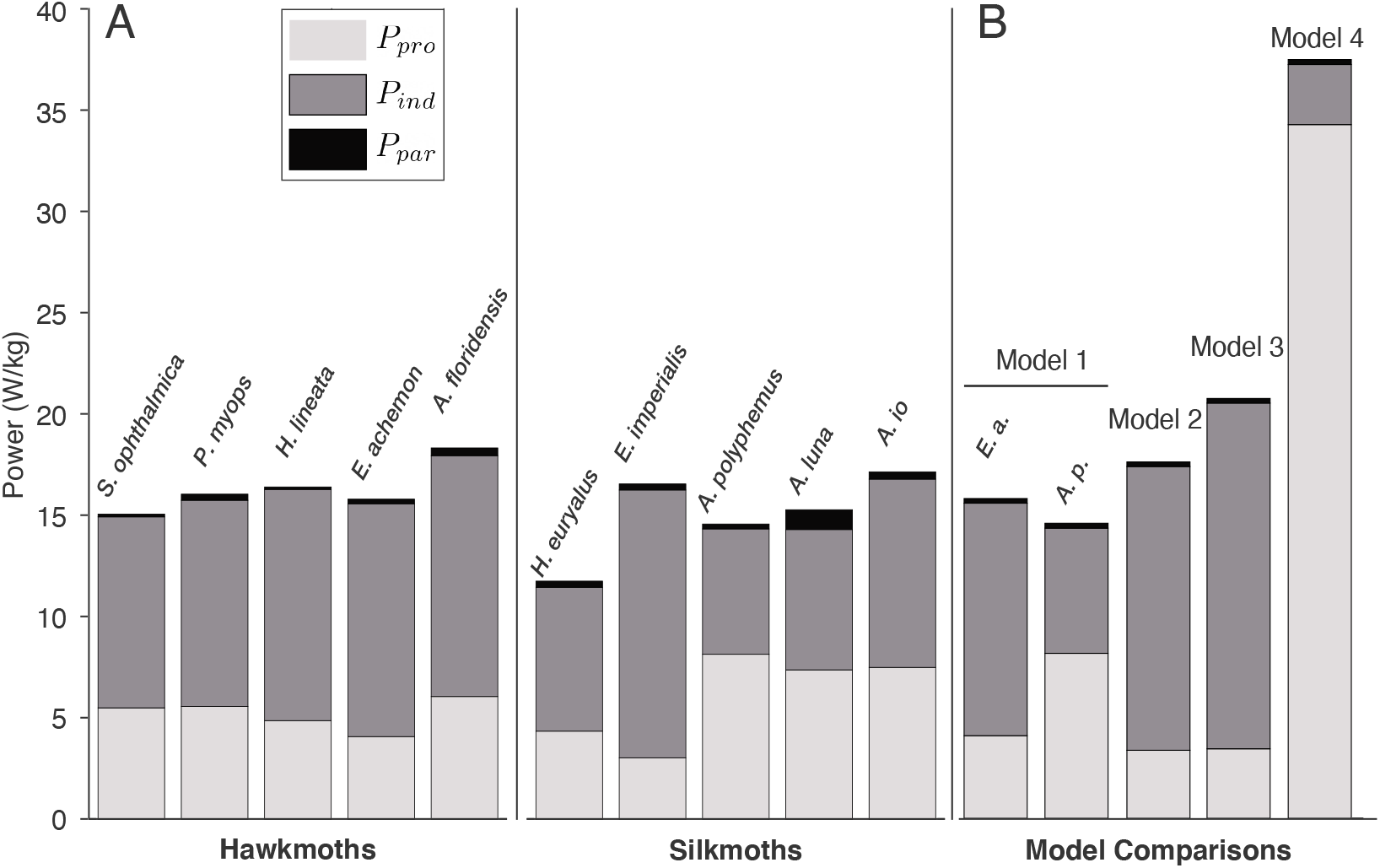
A comparison of the total aerodynamic power requirements of flight. Total aerodynamic power is the sum of induced power (*P*_ind_), profile power (*P*_pro_), and parasitic power (*P*_par_). The total and component power is shown for (A) all species during forward flight at recorded speeds and (B) each sub-model of our two species comparison in Fig. 3 B-D.

### 3.4. Hawkmoths and silkmoths evolved interspecific differences in within-wingstroke aerodynamics

While the average powers and normalized forces converge, the within-wingstroke force profiles do show consistent differences between clades reflecting different strategies for achieving body weight support and thrust. In hawkmoths, 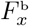 is predominately negative (backwards) during the first half and positive during the second half of the wingstroke. In silkmoths, 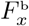 has significant positive and negative portions during both halves of the wing stroke (Figs. 3 A, S6). In hawkmoths, 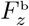 generally supports weight (negative) across the entire wingstroke and of greater (more than twice) magnitude during the first half than the second wingstroke half (Figs. 3 A, S6). In silkmoths, 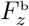 is also generally negative, but usually of reduced magnitude during the second half stroke in comparison to hawkmoths which, in two silkmoth species, becomes predominantly positive (Figs. 3 A, S6). Although the magnitude of rotational 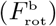 and added mass 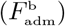 forces are generally larger in silkmoths than hawkmoths (Fig. S6, S7), these forces tend to act in opposition, and, across all species, interspecific differences in total force 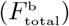 are primarily due to translational force, 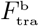 (Fig. S6, S7). The silkmoth *A. io* represents an exception to this as 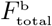 is indeed impacted by 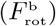 and 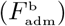 throughout the wingstroke (Fig. S6). These results are consistent whether we consider the species-specific flight speeds or model all species at 2 m s^−1^ (Fig. S8).

### 3.5. Quasi-steady blade element models separate the contributions of wing shape, size, and kinematics to flight strategies

We created several intermediate models to separate contributions and assess how size, shape and kinematics impact aerodynamics. To do so, we focused on exemplar species from each clade (*E. achemon*, *A. polyphemus*), which are representative of the general divergence in wing shape, size, and movement between the clades. The base comparison reported above (Model 1 – Figs. 3 A, and S7A-B) uses each species’ own wing shape and size, time-varying kinematics (*φ*, *θ*, *α*, *β*, *χ*, and *n*), and recorded forward flight velocities. In Model 2 (Figs. 3 B, S7C) kinematics and size are set to that of the hawkmoth *E. achemon* in both cases, leaving interspecific differences only in wing shape. In Model 3 (Figs. 3 C, S7D), wing shape and size are set to that of *E. achemon*, leaving interspecific differences only in wing kinematics. Finally, in Model 4 (Figs. 3 D, S7E), wing shape and kinematics are set to that of *E. achemon*, leaving interspecific differences only in wing size.

#### Wing shape

The *E. achemon* wing shape produces slightly larger net aerodynamic force than *A. polyphemus* shaped wings (Model 2, Figs. 3B, S7C). The increase in 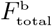 is primarily driven by the increased 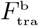 (Fig. S7C). While different in average and peak magnitude, the pattern of the 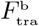 (as well as 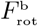) and 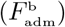 throughout the wing stroke was similar in both wing shapes (Fig. S7C). The *E. achemon* wing shape produced larger 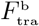 because of its larger radius of second moment of area than *A. polyphemus*.

#### Wing kinematics

The most apparent aerodynamic interspecific difference due to kinematics alone (Model 3) is that *A. polyphemus* kinematics produces lower force (5 times smaller peak force) than *E. achemon* kinematics (Fig. 3C, S7D). The reduction in 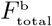 is again determined primarily by differences in 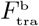 (Fig. S7D). The main cause of this difference is that *n*, and hence wing angular velocity, of *E. achemon* is ~3 times greater than *A. polyphemus* (Fig. 2; Table S3). Interspecific differences in kinematics are also responsible for interspecific differences in the sign of 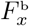 and 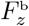 in Model 1 during the first half and second half of wingstroke, respectively (Fig. 3A). To break this down further, we separated the contributions of stroke plane angle (*β*), wing angles (*ϕ*, *θ*, and *α*) and *n* (Fig. S9). The interspecific sign flip in 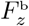 (Fig. 3A) occurs due to *n* (Fig. S9B), and the 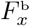 sign flip (Fig. 3A) occurs primarily from differences in lower *n* and more vertical *β* of *A. polyphemus* (Fig. S9B,C).

#### Wing size

*A. polyphemus* has 2.5 times larger wings and, if all other variables are equal (Model 4), it is not surprising that *A. polyphemus* wing size produces about 6 times larger forces (Figs. 3D, S7E). As before, differences in 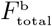 are primarily due to differences in 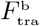 (Fig. S7E). However, the 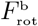) and (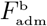 components of 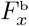 and 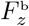 are also nearly an order of magnitude larger in *A. polyphemus* sized wings compared to *E. achemon* sized wings (Fig. S7E). Overall, wing size has the predictable effect of scaling force components.

## 4. Discussion

### 4.1. The evolution of two distinct strategies for flapping flight

Silkmoths and hawkmoths have evolved two distinct strategies (combinations of wing shape, size, and movement) for flapping flight (Fig. 5). Species of both families have evolved sufficient average force production and power reduction, but do so with distinctly different wing shapes, sizes, and movements and different patterns of within-wingstroke force production. An adaptive shift in hawkmoth wing size and shape resulted in the divergence of wing morphology between the two clades [7]. In the present study, we now demonstrate that key wing movement features also diverge between clades, indicating that hawkmoth wing movements, which occur at frequencies more than 3 times greater than silkmoths (Fig. 2), might also be evolving around an adaptive peak.

**Figure 5:**
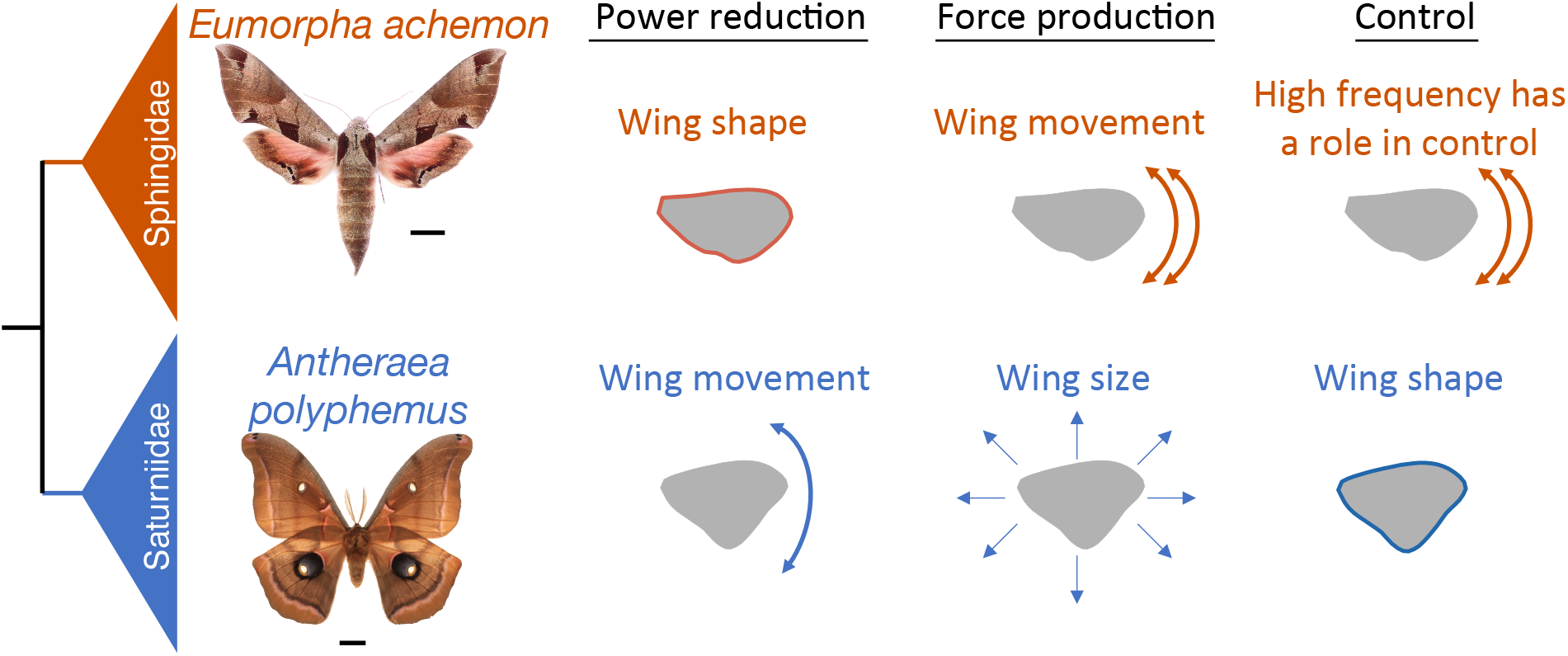
The evolution of two distinct strategies for flight. A summary of how distinct divergence in wing shape, size, and movement impact flight performance in each clade.

In hawkmoths, sufficient force production is primarily due to the evolution of high *n* (movement) (Fig. 2) because translational force production, the largest contributor to total force, is proportional to velocity squared [27]. The evolution of higher 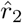 wings (shifting more area distally) in hawkmoths also enhances the force and torque production (e.g. [35, 36]) compared to silkmoths. Silkmoths achieve sufficient force production through relatively larger wings than hawkmoths (Fig. 1 D) because wing area is proportional to translational force [27]. So, hawkmoths primarily use kinematics to enhance the force production, whereas silkmoths primarily use wing shape and size.

Both clades are also likely under selective pressures to reduce the power requirements of flight. Hawkmoths regularly sustain long duration bouts of hovering, often associated with nectaring from flowers [19, 21, 22, 23], which requires a high-power output [28], while all silkmoths do not feed as adults, relying on limited energy reserves gathered as larvae [16, 17]. Hawkmoths evolved high AR wings (shape) ([7]; Fig. 1C), reducing the induced power (*P*_ind_) requirements of flight [37, 38] in comparison the evolution of low AR in silkmoths (Fig. 4B). Silkmoths evolved high amplitude wing strokes (*ϕ*_p-p_), also reducing *P*_ind_ (Fig. 4B), which is inversely proportional to both wing size (*R*) and (*ϕ*_p-p_) [27, 29]. In total, silkmoths have a lower *P*_ind_ than hawkmoths (Fig. 4A), offsetting the larger profile power (*P*_pro_) incurred by the evolution of larger wings in silkmoths than hawkmoths, which ultimately results in similar overall power requirements between the clades (Fig. 4A). Thus, the evolution of hawkmoth wing shape and silkmoth wing movement are different means to reduce power.

Aerodynamic performance, therefore, emerges from the interaction of wing shape, size, and kinematics, demonstrating an example of correlated evolution between the components of a complex locomotor system [39]. The ability for natural selection to act both on wing shape and kinematics to impact force generation and the power requirements of an animal demonstrates the potential decoupling of animal locomotor performance metrics. Because wing shape, size, and movement can each independent affect aerodynamic performance, there are likely fewer purely morphological constraints forcing convergence. This may contribute the diversity of wing shapes across extant aerial animals, even in closely related sister clades.

### 4.2. The evolution of two distinct aerodynamic strategies contributes to inter-clade differences in flight behavior

#### Large, slow wings contribute to bobbing flight behavior in silkmoths

While the two strategies contribute comparable average aerodynamics forces and power, inter-clade differences in within-wingstroke aerodynamics (Fig. 3A) likely contribute to differences in flight behavior between the clades. Silkmoths often display erratic flight patterns [17, 24, 16, 25] where vertical position is regularly changing throughout their flight bout. While all species produce sufficient 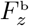 for forward flight, silkmoths generally have more variation in 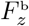 between the first and second half of the wingstroke in comparison to hawkmoths (Fig. 3A, S6). This is especially noticeable in *A. polyphemus* and *A. luna* where 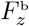 switches sign to become positive positive during the second half of wingstroke. Large force fluctuations and asymmetry (Figs. 3A, S6) do lead to greater fluctuations in body vertical velocity (Figs. S4, S5; Table S3) and are likely the source of the bobbing. Such erratic motions in silkmoths and may be useful in predator avoidance and may limit power minimization despite fixed energy budgets as adults.

#### Evolution of high wing beat frequency (n) could contribute to hawkmoth maneuverability

The divergence of *n* parallels the divergence of flight behavior [16, 17, 18] and wing morphology [7] between hawkmoths and silkmoths. The evolution of high *n* might be key to conducting high speed maneuvers in small flapping flyers like hawkmoths. While we do not directly measure maneuverability in this study, it is clear that hawkmoths have evolved a means to accomplish rapid maneuvers while foraging [19, 22, 23]. As opposed to fixed-wing cases, maneuverability of flapping flight relies on the generation of aerodynamic forces from wing movement to initiate directional change [40]. Therefore, an increase in *n* would allow for more frequent modification of force vectors, which could increase maneuverability. Further, increasing *n* will also enhance maneuverability by increasing the force and torque produced by a wing [41], which is exemplified in Model 3 of this study (Fig. 3C). The diversification of *n*, which supports our previous estimates based on scaling relationships [7], could contribute to interspecific variation in flight control and maneuverability across species. Therefore, we suggest that high n is one of the aspects of flight control that evolved in hawkmoths allowing for the completion of high frequency maneuvers.

### 4.3. Hawkmoth and silkmoth aerodynamics are more similar to other flying insects than they are to each other

Despite being sister clades, the respective flight strategies of hawkmoths and silkmoths are more similar to other flying insects than they are to each other. The hawkmoth flight strategy is most similar to that of other high frequency flappers like flies. In both hawkmoths (Figs. 3A, S6) and the fly, *Drosophila melanogaster*, vertical force is produced during both halves of the wing stroke [42, 43]. The low frequency flapping and time varying aerodynamics of silkmoths are most similar to those of butterflies. Across multiple species of butterflies and silkmoths, vertical force produced during the downstrokes acts to offset body weight while vertical force during the upstroke contributes either minimal or negative body weight support (Fig. 3A); [8]). As silkmoths and butterflies converged in wing shape [6], relative size [6], kinematics [8], and aerodynamics [8], it is not surprising that many species of both groups also evolved erratic flight. Similarly, species of both clades also evolved similar aerodynamic power requirements [8]. As butterflies feed as adults and silkmoths do not, the convergent power requirements suggest silkmoths have not evolved any adaptations for additional power reduction in comparison to butterflies. The combination of wing morphology and kinematics that have evolved in both silkmoths and butterflies might be constrained for this style of flight. Finally, bumblebee aerodynamics at slow flight speeds are similar to that of hawkmoths, but, at fast flight speeds, vertical force production is asymmetric between first and second wing stroke halves [44], similar to a silkmoth (Fig. 3A). The shift in bee aerodynamics is due to an increasingly more vertical stroke plane at greater flight speeds (similar to silkmoths) [45], further exemplifying how the evolution of divergent kinematics between hawkmoths and silkmoths contributes to inter-clade differences in aerodynamics and flight behavior.

In conclusion, the divergence in flight strategy between the hawkmoth and silkmoth sister clades is just as strong as the evolutionary divergence between distant clades. Through a strong divergence in wing shape, size, and movement, hawkmoths and silkmoths have evolved two independent flight strategies. This study thus demonstrates that even closely related clades with recently shared ancestry can still undergo strong divergence in flight strategies. The change in wing size and shape corresponds to a adaptive shift between the silkmoths and hawkmoths [7], which is now reflected in their kinematics and performance. These sister groups are ideal “model clades” [46] for exploring the morphological, neuromechanical, and behavior diversification of insect flight. Finally, while the design of human engineered flapping fliers is often attributed to inspiration by insect flight, a rich design space might be better inspired by the many strategies that even closely related insects take to achieve similar but flexible flight behaviors.

## Supporting information

Supplemental methods and results

## ACKNOWLEDGMENTS

We thank Jesse Barber, Stephanie Gage, Jeff Gau, Megan Matthews, Izaak Neveln, David Plotkin, Joy Putney, Juliette Rubin, Varun Sharma, Ryan St Laurent, and Travis Tune for helpful discussions, Aaron Olsen for assistance with image digitization and analysis, Bo Cheng and Yagiz Bayiz for advice on the aerodynamic blade element model, and Laurel Kaminsky for assistance in imaging museum specimens.

## References

[1] D. B. Nicholson, A. J. Ross, P. J. Mayhew, Fossil evidence for key innovations in the evolution of insect diversity, Proceedings of the Royal Society B: Biological Sciences 281 (1793) (2014) 20141823.

[2] D. R. Warrick, K. P. Dial, Kinematic, aerodynamic and anatomical mechanisms in the slow, maneuvering flight of pigeons, Journal of Experimental Biology 201 (5) (1998) 655–672.

[3] S. P. Sane, The aerodynamics of insect flight, Journal of experimental biology 206 (23) (2003) 4191–4208.

[4] R. J. Wootton, Functional-Morphology of Insect Wings, Annual Review of Entomology 37 (1992) 113–140. doi:Doi 10.1146/Annurev.En.37.010192.000553.

[5] S. Combes, T. Daniel, Flexural stiffness in insect wings i. scaling and the influence of wing venation, Journal of experimental biology 206 (17) (2003) 2979–2987.

[6] C. Le Roy, V. Debat, V. Llaurens, Adaptive evolution of butterfly wing shape: from morphology to behaviour, Biological Reviews 94 (4) (2019) 1261–1281. doi:10.1111/brv.12500.

[7] B. R. Aiello, M. Tan, U. Bin Sikandar, A. J. Alvey, B. Bhinderwala, K. Kimball, J. Barber, C. A. Hamilton, A. Y. Kawahara, S. Sponberg, Adaptive shifts underlie the divergence in wing morphology in bombycoid moths, bioRxiv (2021). arXiv: https://www.biorxiv.org/content/early/2021/06/24/2021.06.23.449655.full.pdf, doi:10.1101/2021.06.23.449655.

[8] R. Dudley, Biomechanics of flight in neotropical butterflies: aerodynamics and mechanical power requirements, Journal of Experimental Biology 159 (1) (1991) 335–357.

[9] Y. H. J. Fei, J. T. Yang, Importance of body rotation during the flight of a butterfly, Physical Review E 93 (3) (2016) 1–10. doi:10.1103/PhysRevE.93.033124.

[10] S. P. Sane, M. H. Dickinson, The aerodynamic effects of wing rotation and a revised quasi-steady model of flapping flight, Journal of Experimental Biology 205 (8) (2002) 1087–1096.

[11] J. S. Han, J. K. Kim, J. W. Chang, J. H. Han, An improved quasi-steady aerodynamic model for insect wings that considers movement of the center of pressure, Bioinspiration and Biomimetics 10 (4) (2015). doi:10.1088/1748-3190/10/4/046014.

[12] R. J. Bomphrey, T. Nakata, N. Phillips, S. M. Walker, Smart wing rotation and trailing-edge vortices enable high frequency mosquito flight, Nature 544 (7648) (2017) 92–95.

[13] J. K. Wang, M. Sun, A computational study of the aerodynamics and forewing-hindwing interaction of a model dragonfly in forward flight, Journal of experimental biology 208 (19) (2005) 3785–3804.

[14] A. Y. Kawahara, D. Plotkin, M. Espeland, K. Meusemann, E. F. A. Toussaint, A. Donath, F. Gimnich, P. B. Frandsen, A. Zwick, M. dos Reis, J. R. Barber, R. S. Peters, S. L. Liu, X. Zhou, C. Mayer, L. Podsiadlowski, C. Storer, J. E. Yack, B. Misof, J. W. Breinholt, Phylogenomics reveals the evolutionary timing and pattern of butterflies and moths, Proc Natl Acad Sci USA 116 (45) (2019) 22657–22663. doi:10.1073/pnas.1907847116.

[15] C. A. Hamilton, R. A. St Laurent, K. Dexter, I. J. Kitching, J. W. Breinholt, A. Zwick, M. Timmermans, J. R. Barber, A. Y. Kawahara, Phylogenomics resolves major relationships and reveals significant diversification rate shifts in the evolution of silk moths and relatives, BMC Evol Biol 19 (1) (2019) 182. doi:10.1186/s12862-019-1505-1.

[16] D. H. Janzen, Two ways to be a tropical big moth: Santa Rosa saturniids and sphingids, in: D. R. R. M (Ed.), Oxford surveys in evolutionary biology, Oxford University Press, Oxford, 1984, pp. 85–140.

[17] P. M. Tuskes, J. P. Tuttle, M. M. Collins, The Wild Silk Moths of North America: A Natural History of the Saturniidae of the United States and Canada, Cornell University Press, Ithica, NY, 1996.

[18] J. Tuttle, The Hawk Moths of North America: A Natural History Study of the Sphingidae of the United States and Canada (2007).

[19] L. Wasserthal, Swing-hovering combined with long tongue in hawkmoths, an antipredator adaptation during flower visits., in: B. W. N. C. S.-L. K. S. LL (Ed.), Animal-plant interactions in tropical environments., Zoologisches Forschungsinstitut und Museum König, Bonn, 1993, pp. 77–87.

[20] W. M. Farina, D. Varju, Y. Zhou, The Regulation of Distance to Dummy Flowers during Hovering Flight in the Hawk Moth Macroglossum-Stellatarum, Journal of Comparative Physiology a-Sensory Neural and Behavioral Physiology 174 (2) (1994) 239–247.

[21] J. D. H. Sprayberry, T. L. Daniel, Flower tracking in hawkmoths: behavior and energetics, Journal of Experimental Biology 210 (1) (2007) 37–45. doi:10.1242/jeb.02616.

[22] S. Sponberg, J. P. Dyhr, R. W. Hall, T. L. Daniel, Luminance-dependent visual processing enables moth flight in low light, Science 348 (6240) (2015) 1245–1248.

[23] A. L. Stoöckl, K. Kihlstroöm, S. Chandler, S. Sponberg, Comparative system identification of flower tracking performance in three hawkmoth species reveals adaptations for dim light vision, Philosophical Transactions of the Royal Society B-Biological Sciences 372 (1717) (2017) 20160078–20160079.

[24] D. S. Jacobs, A. Bastian, Predator-prey interactions: co-evolution between bats and their prey., Springer, Cham, Switzerland, 2016.

[25] F. P. Lewis, J. H. Fullard, S. B. Morrill, Auditory Influences on the Flight Behavior of Moths in a Nearctic Site .2. Flight Times, Heights, and Erraticism, Canadian Journal of Zoology-Revue Canadienne De Zoologie 71 (8) (1993) 1562–1568. doi:DOI 10.1139/z93-221.

[26] B. Cheng, B. W. Tobalske, D. R. Powers, T. L. Hedrick, Y. Wang, S. M. Wethington, G. T. C. Chiu, X. Y. Deng, Flight mechanics and control of escape manoeuvres in hummingbirds. II. Aerodynamic force production, flight control and performance limitations, Journal of Experimental Biology 219 (22) (2016) 3532–3543. doi:10.1242/jeb.137570.

[27] C. P. Ellington, The Aerodynamics of Hovering Insect Flight 6. Lift and Power Requirements, Philosophical Transactions of the Royal Society of London Series B-Biological Sciences 305 (1122) (1984) 145–181. doi:Doi 10.1098/Rstb.1984.0054.

[28] A. P. Willmott, C. P. Ellington, The mechanics of flight in the hawkmoth *Manduca sexta*. II. Aerodynamic consequences of kinematic and morphological variation., Journal of Experimental Biology 200 (21) (1997) 2723–2745.

[29] K. Warfvinge, M. KleinHeerenbrink, A. Hedenstrom, The power-speed relationship is U-shaped in two free-flying hawkmoths (Manduca sexta), Journal of the Royal Society Interface 14 (134) (2017). doi:ARTN 2017037210.1098/rsif.2017.0372.

[30] A. M. Olsen, M. W. Westneat, StereoMorph: an R package for the collection of 3D landmarks and curves using a stereo camera set-up, Methods in Ecology and Evolution 6 (3) (2015) 351–356. doi:10.1111/2041-210X.12326.

[31] C. P. Ellington, The Aerodynamics of Hovering Insect Flight .2. Morphological Parameters, Philosophical Transactions of the Royal Society of London Series B-Biological Sciences 305 (1122) (1984) 17–40. doi: Doi 10.1098/Rstb.1984.0050.

[32] M. Matthews, S. Sponberg, Hawkmoth flight in the unsteady wakes of flowers, The Journal of Experimental Biology 221 (22) (2018) jeb179259. doi:10.1242/jeb.179259.

[33] B. J. Knorlein, D. B. Baier, S. M. Gatesy, J. D. Laurence-Chasen, E. L. Brainerd, Validation of XMALab software for marker-based XROMM, Journal of Experimental Biology 219 (23) (2016) 3701–3711. doi:10.1242/jeb.145383.

[34] J.-K. Kim, J.-S. Han, J.-S. Lee, J.-H. Han, Hovering and forward flight of the hawkmoth *Manduca sexta*: trim search and 6-DOF dynamic stability characterization, Bioinspiration & Biomimetics 10 (5) (2015) 56012. doi:10.1088/1748-3190/10/5/056012.

[35] F. T. Muijres, N. A. Iwasaki, M. J. Elzinga, J. M. Melis, M. H. Dickinson, Flies compensate for unilateral wing damage through modular adjustments of wing and body kinematics, Interface Focus 7 (1) (2017). doi:ARTN 2016010310.1098/rsfs.2016.0103.

[36] M. J. Fernandez, M. E. Driver, T. L. Hedrick, Asymmetry costs: effects of wing damage on hovering flight performance in the hawkmoth Manduca sexta, Journal of Experimental Biology 220 (20) (2017) 3649–3656. doi:10.1242/jeb.153494.

[37] U. M. Norberg, J. M. V. Rayner, Ecological Morphology and Flight in Bats (Mammalia, Chiroptera) - Wing Adaptations, Flight Performance, Foraging Strategy and Echolocation, Philosophical Transactions of the Royal Society B-Biological Sciences 316 (1179) (1987) 337–419.

[38] C. J. Pennycuick, Power Requirements for Horizontal Flight in Pigeon Columba Livia, Journal of Experimental Biology 49 (3) (1968) 527–+.

[39] B. R. Aiello, M. W. Westneat, M. E. Hale, Mechanosensation is evolutionarily tuned to locomotor mechanics, Proc Natl Acad Sci USA 14 (17) (2017) 4459–4464. doi:10.1073/pnas.1616839114.

[40] D. R. Warrick, K. P. Dial, A. A. Biewener, Asymmetrical force production in the slow maneuvering flight of pigeons, Auk 115 (1998) 916–928.

[41] T. L. Hedrick, B. Cheng, X. Deng, Wingbeat time and the scaling of passive rotational damping in flapping flight, Science 324 (5924) (2009) 252–255. doi:10.1126/science.1168431.

[42] S. N. Fry, R. Sayaman, M. H. Dickinson, The aerodynamics of hovering flight in drosophila, Journal of Experimental Biology 208 (12) (2005) 2303–2318.

[43] M. H. Dickinson, F. T. Muijres, The aerodynamics and control of free flight manoeuvres in drosophila, Philosophical Transactions of the Royal Society B: Biological Sciences 371 (1704) (2016) 20150388.

[44] R. Dudley, C. Ellington, Mechanics of forward flight in bumblebees: Ii. quasi-steady lift and power requirements, Journal of Experimental Biology 148 (1) (1990) 53–88.

[45] R. Dudley, C. Ellington, Mechanics of forward flight in bumblebees: I. kinematics and morphology, Journal of Experimental Biology 148 (1) (1990) 19–52.

[46] N. Jourjine, H. E. Hoekstra, Expanding evolutionary neuroscience: insights from comparing variation in behavior, Neuron (2021).

