## Supplemental methods and results for "The evolution of two distinct strategies of moth flight"

#### This PDF file includes:

Supplementary text
Figs. S1 to S9
Tables S1 to S4
References for SI reference citations

### Supporting Information Text

### 1. Materials and methods

Definitions and details of the mathematical notation and symbols used throughout this study are in Table S1.

**A. Live specimens.** Live specimens of five different species from each sister-family (10 total species) were used in this study (Table S2). Species from the hawkmoth (Sphingidae) family include: *Eumorpha achemon*, *Amphion floridensis*, *Hyles lineata*, *Paonias myops*, and *Smerinthus ophthalmica*. Species from the silkmoth (Saturniidae) family include: *Actias luna*, *Automeris io*, *Antheraea polyphemus*, *Hyalophora euryalus*, and *Eacles imperialis*. The species from each family were chosen because they were locally available in large numbers, provide a sufficient representation of the variation in wing morphology between and within each family (1), and provide a generally even distribution across the phylogeny (i.e. they are not all clustered in the phylogeny). Caterpillars of each species were acquired by collecting eggs from local adult moths, and all caterpillars were reared on species-specific host plants. Pupae were stored in an incubator (Darwin Chambers, model: IN034LTDMMP, Saint Louis, MO) set to a temperature of 23° C and a relative humidity of 65%.

**B. Body and Wing Measurements and Morphometrics.** The body and wing morphology was digitized for each live specimen using the StereoMorph package (version 1.6.2) (2) in R (version 3.4.2; The R Foundation for Statistical Computing). Body and wing landmarks follow our previous methodology (1). For all individuals, total body mass ($m_{\text{t}}$) was measured directly after the individual was flown in the wind tunnel. Forewing and hindwing masses ($m_{\text{fw}}$, $m_{\text{hw}}$) was estimated from our previously established scaling relationships between wing mass and area of this moth clade (1) and further confirmed with 5 individuals of this study.

Morphology was analyzed in MATLAB (version R2018b - 9.5.0.944444), following (1). To generate a combined wing shape from the overlap of the fore- and hindwings, the forewing was rotated so its long axis was perpendicular to the long axis of the body. In hawkmoths, the long axis of the hindwind was also oriented perpendicular to the long axis of the body. In silkmoths, the orientation of the hindwing was left in its natural position obtained when the wings are splayed open and the moth is at rest. This position was chosen because while reviewing videos of silkmoth flight, the long axis of hindwing is always oriented posteriorly and nearly parallel to the body long axis (see supplemental videos). All wing morphology parameters (see supplement) needed for the blade element model were calculated from the combined wing and follow (3).

**C. High speed recordings of moth flight.** Moths were transferred to the wind tunnel in individual containers with a moist tissue to prevent desiccation. Other than individuals of *A. floridensis*, which is a diurnal species, each individual was dark adapted at the wind tunnel for 1hr prior to the start of filming. Flight experiments were conducted in a 100×60.96 working section of an open-circuit Eiffel-type wind tunnel (ELD, Inc, Lake City, MN). The stream-wise turbulence of the wind tunnel does not exceed 0.5% and the flow speed did not vary by more than 2%. For a detailed overview of the specifications of the wind tunnel see (4).

Moths were enticed to fly by providing a mild wind speed of 0.7 ms$^{-1}$. Flight bouts were filmed at 2000 frames s$^{-1}$ for hawkmoths and 1000 frames s$^{-1}$ for silkmoths using three synchronized Photron high-speed digital video cameras (Mini UX 100; Photron, San Diego, CA, USA) at a resolution of 1280×1024. Two cameras (one upwind and one downwind) were positioned below the wind tunnel test section at a 45° angle relative to the direction of flow. A third camera was placed laterally and orthogonal to plane of the first two cameras. The working section of the wind tunnel was illuminated with six 850Nm IR lights (Larson Electronics, Kemp, TX, USA) and a neutral density filter, white LED "moon" light (Neewer CW-126) to control illumination conditions (5). For the diurnal species (*A. floridensis*), the room lights were also turned on.

**D. Extracting the 3D Time-series of Moth Wing and Body Motion.** Videos were digitized and calibrated in XMALab (6). A total of seven landmarks were digitized on the moth: rostral tip of the head (between the antennae), junction between the thorax and abdomen, caudal tip of the abdomen, left and right forewing hinges, right wing tip, and a point on the trailing edge of the wing. The coordinates of head, thorax and abdomen were tracked to determine the orientation of the body. The points on the right wing hinge, right wing tip and trailing edge were used to determine the wing kinematics. We only extracted data from forward flight bouts, avoiding initial and final wingstrokes. From each individual, we digitized at least 1 complete wingstroke that was as close to steady forward flight as possible and contained within a larger set of wingstrokes during forward flight (See Table S3 for the number of wingstrokes captured for each individual). For each wing stroke, at minimum, every other frame was digitized, and, in most videos, we digitized every frame. The time-varying trajectories of each landmark were linearly interpolated for any frame that was not digitized and then smoothed using a moving-average filter with a window length of 10 frames.

**E. Blade Element model summary.** Extracted time series of 3D trajectories of body and wings were used to compute time-varying body and wing kinematic angles and body speeds. These time-varying parameters were then separated into wingstrokes. A large majority of the waveforms of these parameters were roughly periodic, so a third-order Fourier series was fitted to each these kinematic parameters for every wingstroke. Winsgrokes with at least one non-periodic parameter waveform were excluded from the data to ensure the consistency of the steady flight assumption. Then, for each species, we averaged the wing shapes and time-varying Fourier-fitted kinematics over all wingstrokes across all individual moths to obtain representative wingstroke kinematics for each species.

The species-specific aerodynamic forces were evaluated using a quasi-steady blade element model. We assumed a wing as a thin rigid plate divided into 200 chord-wise strips, where each strip is treated as an independent airfoil. Briefly, the model estimated the total aerodynamic force on a strip as contributions from forces due to translational and rotational motion of the strip, and the force due to added mass (7–12). The lift and drag coefficients were based on empirical measurements from dynamically scaled wings of hawkmoth *Manduca sexta*. Similarly measured coefficients were not available for the species of moths used in this study. However, the fact that our aerodynamic model was able to achieve force and moment equilibrium qualifies this as a good assumption.

We first calculated all aerodynamics in a wing-attached coordinate frame and summed across all strips to calculate the total force on the wing. Then the force was transformed to the body-attached frame to determine its vertical and fore-aft components. To find the force and moment equilibrium over a wingstroke, the aerodynamic model was trimmed to make the vertical force balance weight and cancel out the fore-aft force and pitching moment. The trim search was performed in the mean and amplitude space of Fourier-fitted kinematics of body and wings. The trim search space was restricted to the minimum and maximum values of mean and amplitude observed for each species in our data. After the trim search, the mean and amplitude of each species-specific time-varying kinematic parameter were adjusted to the values found in the trim search. The readjusted kinematics were used to calculate the aerodynamic forces from the blade-element model. We also calculated the total aerodynamic power as a sum of induced, profile and parasite powers (13). A detailed formulation of each step of the blade element model can be found in subsequent sections.

In our kinematic and aerodynamic model formulation, we built up on but made a few considerable modifications to the previously used methods (8, 12, 14, 15). We used fully time-varying stroke-plane angle, body pitch angle and body velocity, which is rarely found in the previous models. In addition to including a stroke-plane angle which is a title about a lateral axis, we also included a small stroke-plane tilt about the anterior-posterior axis because it was measured in our kinematic data. But its value was assumed constant over a wingstroke. Regarding trimming the aerodynamic model, previously, the natural limitations of an animal's anatomical and kinematic capabilities have not been usually considered (9, 11). But in our trim search, we restricted our parameter search space between the species-specific maximum and minimum values that we observed in our data. Lastly, in our pitching moment calculation, we also included the moment arm between the body center of mass and the wing hinge.

To explore the aerodynamic effects of morphological and kinematic parameters, we assumed different configurations of the model to assess the relative contribution of wing shape, size and kinematics to the aerodynamic force and power production.

**F. Coordinate Frames of the Moth Wing and Body.** We defined four coordinate frames to track the 3-D position and orientation of the moth's body, calculate the wing kinematic angles, and translate the aerodynamic forces from the wing frame to the body frame.

***F.1. Body Coordinate Frames.*** A body-attached frame specifies the direction of flight relative to the absolute horizontal and a body-long frame specifies the body's 3-D orientation (Fig. S1A). Both frames share a common origin at the center of mass. The body-long positive $x$-axis, $x^l$, points from the center of mass towards the center of head; the $z^l$-axis points ventrally and lies in the vertical saggital plane, which splits the moth body into bilaterally symmetric halves; the $y^l$-axis is the cross product of $z^l$ and $x^l$ according to a right-handed coordinate system. The body-attached positive $x$-axis, $x^{\mathrm{b}}$, starts from the center of mass and points in the direction of the $x^l$-axis projection on the absolute horizontal plane; $z^{\mathrm{b}}$ points in the direction of gravity; $y^{\mathrm{b}}$ is the cross product of $z^{\mathrm{b}}$ and $x^{\mathrm{b}}$, making the $x^{\mathrm{b}}y^{\mathrm{b}}$-plane the absolute horizontal plane irrespective of body orientation. In other words, the body-attached frame is invariant under pitch and roll rotations of the body. The direction of gravity was determined using a plumbline hung in the working section of the wind tunnel after each recording.

***F.2. Stroke-plane Coordinate Frame of the Right Wing.*** The origin of a stroke-plane frame is at the wing hinge point as shown in Fig. S1A . Anatomically, we defined the wing hinge point as a single point located at one-third the distance from the rostral to the caudal wing hinge. For the right wing, positive $x^{\mathrm{s}}$-axis is in the direction of the downstroke and lies within the $x^{\mathrm{b}}$-$z^{\mathrm{b}}$ plane; $y^{\mathrm{s}}$ is outward from the right wing hinge parallel to the $y^l$-axis in a direction from the left wing hinge to the right wing hinge; and $z^{\mathrm{s}}$ is the cross-product of $x^{\mathrm{s}}$ and $y^{\mathrm{s}}$.

***F.3. Wing-attached Coordinate Frame of the Right Wing.*** To calculate the forces on the wing at each time instance, we rely on wing-attached coordinate frame S1D. Origin of this frame is also at the wing hinge point. Its $y$-axis, $y^{\mathrm{w}}$, is the anatomical pitching axis of the wing, which was set to be perpendicular to the body-long axis and lie in the same plane as the wing. Its $x^{\mathrm{w}}$ axis lies in the stroke-plane and $z^{\mathrm{w}}$ is the cross product of $x^{\mathrm{w}}$ and $y^{\mathrm{w}}$. It is important to note that the wing-attached frame rotates with the sweep ($\phi$) and deviation ($\theta$) rotations of the wing but for simplicity it is invariant under the feathering angle ($\alpha$) rotation.

**G. Body and Wing Kinematics.** The exported 3-D points of the landmarks on head, thorax, abdomen and wing were used to characterize the body and right wing kinematics by calculating the following variables: body angle ($\chi$), body velocity (forward $u$, sideslip $v$ and vertical $w$), stroke-plane angle ($\beta$), stroke-plane roll angle ($\beta_{\mathrm{r}}$), wing beat frequency ($n$), sweep angle ($\phi$), deviation angle ($\theta$) and feathering angle ($\alpha$) (Figs. S1 B-E).

Body angle ($\chi$) is the angle that $x^l$-axis makes with the absolute horizontal. Body velocities $u$, $v$ and $w$ are in the direction of $x^{\mathrm{b}}$, $y^{\mathrm{b}}$ and $z^{\mathrm{b}}$ axes respectively. Stroke-plane angle ($\beta$) and stroke-plane roll angle ($\beta_{\mathrm{r}}$) define the pitch and roll angles of the stroke-plane of each wing with respect to the body-attached frame. In addition to pitch angles which are called stroke-plane angles ($\beta$), most of the stroke planes we determined had slightly tilted roll angles with respect to the absolute horizontal. So we also included a stroke-plane roll angle ($\beta_{\mathrm{r}}$) in our kinematic model. Sweep ($\phi$) and deviation ($\theta$) angles respectively, are the azimuth and elevation angles of the wing-tip from the wing-hinge in the stroke-plane frame. Feathering angle ($\alpha$) is the angle that the wing chord makes with the stroke plane.

The extraction of the time series of these kinematic parameters for wingstrokes was performed in a number of sequential steps. First, the raw time series data of 3-D coordinates of landmarks was filtered using a moving-average filter of window size equal to 10. Then body-longitudinal and body-attached axes were calculated and the thorax landmark point was assumed as the center of mass and hence the origin of these axes. The $z$-axis of the body-attached frame, which is in the direction of gravity, was calculated using three landmarks on the plumbline. All landmark points were then transformed from the camera-calibrated frame to the body-longitudinal frame. To remove any jitter in the points that are supposed to remain roughly fixed with respect to the body-longitudinal frame (head and wing hinges), the points were averaged over all frames of a video. Individual wing strokes were then isolated. A wing stroke was defined to start at the onset of the downstroke and to end at the cessation of the subsequent upstroke, which were determined from the waveform of $phi$. The wingbeat frequency ($n$) is the reciprocal of this period. The stroke plane was determined for each wing stroke using a least-squares line through the 3D wing tip trajectory and right wing hinge point (the definition of stroke-plane was similar to (16)). The stroke plane was fit to each wingstroke separately and a stroke-plane axis was specified. Next, we calculated the three angles of the wing kinematics: the wing sweep angle ($\phi$), the deviation angle ($\theta$) and the wing pitching (feathering) angle $\alpha$ as defined in Figs. S1B and S1C. In the final step, all points were transformed to the body-attached frame to calculate the body angle ($\chi$), the stroke-plane angle ($\beta$), the stroke-plane roll angle ($\beta_{\mathrm{r}}$) and the body velocity ($u$, $v$ and $w$), assuming that during one wingstroke the stroke plane did not rotate with respect to the body-longitudinal frame.

**H. Fitting a Fourier Series to the Wing Kinematics.** For each time-varying parameter, namely $\chi$, $\beta$, $u$, $w$, $\phi$, $\theta$ and $\alpha$, we fit a third-order Fourier series in each wingstroke using lsqcurvefit() function in MATLAB, *e.g.*,

$$\phi(t) \quad = a_{\phi,0} + \sum_{k=1}^{3} a_{\phi,k} \cos(2\pi knt) + b_{\phi,k} \sin(2\pi knt). \qquad [1]$$

where $n$ is the wingbeat frequency and $a_{\phi,k}$ and $b_{\phi,k}$ are the Fourier series coefficients. Different from the previous blade-element models, we used time-varying $\chi$, $\beta$, $u$ and $w$ instead of assuming wingstroke-averaged constant values. This is because silkmoths have significant within-wingstroke variation in these parameters–large enough to impact the aerodynamics. Regarding the kinematics that affect the lateral body dynamics, because our data represents steady-state forward flight, time-series of velocities of side-slip, roll and yaw were assumed to be equal to zero because these were negligible.

**I. The blade element model.** For each species, we averaged shapes and fully time-varying kinematics of body and wings over all wingstrokes to calculate the aerodynamic forces. We used a blade element model to evaluate the quasi-steady aerodynamic forces produced during forward flight. We assumed a wing as a thin rigid plate divided into 200 chord-wise strips, where each strip is treated as an independent airfoil. Briefly, the model estimated the total aerodynamic force on a strip as contributions from forces due to translational and rotational motion of the strip, and the force due to added mass (7–12). We first calculated all aerodynamics in a wing-attached coordinate frame and summed across all strips to calculate the total force on the wing. Then the force was transformed to the body-attached frame to determine its vertical and fore-aft components. We also calculated the total aerodynamic power as a sum of induced, profile and parasite powers (13). A detailed formulation of each step of the blade element model can be found in subsequent sections. Details of the mathematical notation used are in Table S1. Finally, we used different configurations of the model to assess the relative contribution of wing shape, size and kinematics to the aerodynamic force production.

***I.1. Relative Airflow Velocity.*** We defined the relative airflow velocity $\boldsymbol{V}$ of a small blade element strip as the velocity of the airflow in the far field relative to the strip. This relative airflow is caused by the motion of the strip relative to the surrounding air due to its rotation about the wing hinge, body translation and rotation, and wind velocity.

$$\boldsymbol{V}^{\mathrm{w}} \quad = - \left( \boldsymbol{V}_{\mathrm{b}}^{\mathrm{w}} + \boldsymbol{\omega}_{\mathrm{b}}^{\mathrm{w}} \times \boldsymbol{l}_{3}^{\mathrm{w}} + \begin{bmatrix} -r\dot{\phi} \\ 0 \\ r\dot{\theta} \end{bmatrix} \right) \qquad [2]$$

where $\boldsymbol{V}_{\mathrm{b}}^{\mathrm{w}}$ is the body velocity relative to the wind but measured in the wing-attached frame, $\boldsymbol{\omega}_{\mathrm{b}}^{\mathrm{w}}$ is the body angular velocity pseudovector in the wing-attached coordinate frame, $\boldsymbol{l}_{3}^{\mathrm{w}}$ is the vector from the body center of mass to the center of the strip, $r$ is the distance from wing hinge to the vertical mid-chord line of the strip (see Fig. S1G), and $\dot{\phi}$ and $\dot{\theta}$ are the stroke positional (sweep) and stroke deviation angular velocities of the wing, respectively. For the calculations performed on the data relevant to this paper, there were no body rotations so $\boldsymbol{\omega}_{\mathrm{b}}^{\mathrm{w}}$ was equal to zero. The induced velocity ($\boldsymbol{V}_{\mathrm{ind}}$) of airflow is not included in the relative airflow velocity ($\boldsymbol{V}$) because the induced velocity acts in the near field.

***I.2. The Angle of Attack.*** The angle of attack of the strip is defined as the angle between the chord line vector from the leading edge to the trailing edge and the relative airflow velocity vector. This angle is calculated as

$$\alpha_{\mathrm{e}} = \cos^{-1}\left(-\hat{\boldsymbol{b}}^{\mathrm{w}} \cdot \hat{\boldsymbol{V}}^{\mathrm{w}}\right), \quad \alpha_e \in [0, \pi). \qquad [3]$$

where $\hat{\boldsymbol{b}}^{\mathrm{w}} = \begin{bmatrix} \cos\alpha & 0 & -\sin\alpha \end{bmatrix}^{\top}$ is the unit vector along the chord line in the direction from the trailing edge of the strip to its leading edge, and $\alpha$ is the feathering angle of the strip. But for lift and drag coefficient calculations, we used a restricted

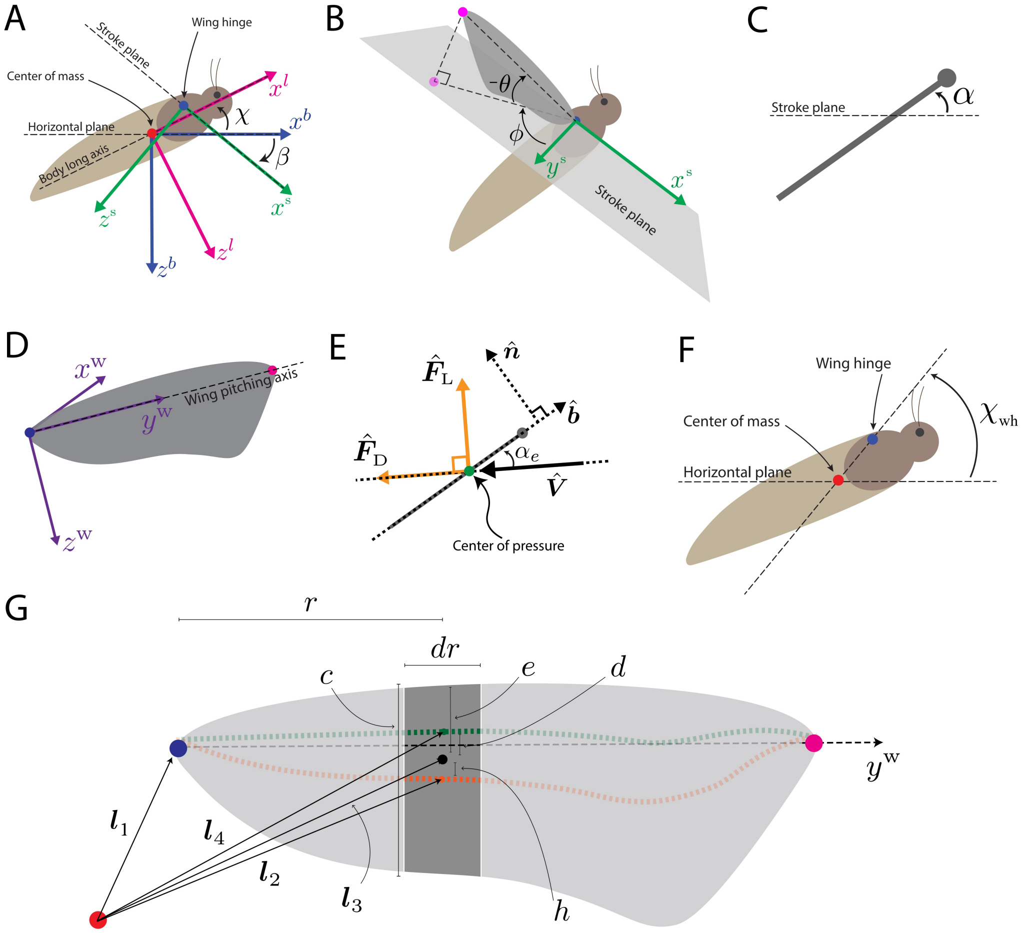


**Fig. S1.** **A** The body-attached coordinate frame (blue), the body-long coordinate frame (pink) and the stroke-plane frame (green). The red dot represents the location of the center of mass and the blue dot represents the wing hinge. $\beta$ is the stroke-plane angle and $\chi$ is the body angle. **B**-**C** Definitions of wing kinematic angles: $\phi$ (sweep), $\theta$ (deviation) and $\alpha$ (feathering) defined with respect to the stroke-plane. **D** The wing-attached coordinate frame. **E** Relative airflow, angle of attack, and lift and drag components of the translational aerodynamic force. **F** Elevation angle $\chi_{\mathrm{wh}}$ of the wing-hinge point (blue) from the center of mass (red). **G** Various length parameters relevant to a single wing strip. Red, blue, and pink circles correspond to the body center of mass, wing hinge, and wing tip. Dashed green, orange, and black lines are the quarter-chord, half-chord, and wing pitching axis, respectively.

angle of attack $\alpha_r$ after setting bounds on the values of the angle of attack $\alpha_e$ so that it remains between 0 and $\pi/2$ radians.

$$\alpha_r = \begin{cases} \alpha_e & 0 \le \alpha_e \le \frac{\pi}{2} \\ \pi - \alpha_e & \frac{\pi}{2} < \alpha_e \le \pi \end{cases} \quad [4]$$

This was done because the coefficients we used from (10) were experimentally measured for the angles of attack only in the range from 0 to $\pi/2$ radians. Moreover, this definition of the angle of attack keeps the lift and drag coefficients positive and simplifies the model because the direction of the lift can be specified by the lift force direction vector $\hat{\boldsymbol{F}}_L$ (see the next section).

***I.3. Translational Aerodynamic Force.*** The translational aerodynamic force is the sum of the lift and drag forces on the wing and acts at the center of pressure. We assumed the center of pressure to be located on the wing at a distance one-quarter chord length behind the leading edge (green dashed line in Fig. S1G), because this is the region at which the bound vortex has been regarded to be concentrated according to the thin airfoil theory for both steady and unsteady aerodynamic effects (17). The lift and drag forces were calculated using the aerodynamic coefficients of hawkmoth *Manduca sexta* taken from (10). The equations of these forces acting on a small wing strip of width $dr$ are as follows (12).

$$d\boldsymbol{F}_L^w = \frac{1}{2}\rho C_L V^2 c \; dr \; \hat{\boldsymbol{F}}_L^w \; , \quad [5]$$

$$d\boldsymbol{F}_D^w = \frac{1}{2}\rho C_D V^2 c \; dr \; \hat{\boldsymbol{F}}_D^w \; , \quad [6]$$

where $\rho$ is the air density, the aerodynamic coefficients (10)

$$C_L(\alpha_r) = 1.552 \sin\alpha_r \cos\alpha_r + 1.725 \sin^2\alpha_r \cos\alpha_r, \quad [7]$$

$$C_D(\alpha_r) = 0.0596 \sin\alpha_r \cos\alpha_r + 3.598 \sin^3\alpha_r, \quad [8]$$

$V$ is the relative airflow speed of the strip, $c$ and $dr$ are chord length and width of a strip, and the translational drag and lift unit vectors, $\hat{\boldsymbol{F}}_L$ and $\hat{\boldsymbol{F}}_D$, are calculated as follows

$$\hat{\boldsymbol{F}}_L^w = \frac{\boldsymbol{q}^w}{|\boldsymbol{q}^w|}, \quad [9]$$

$$\hat{\boldsymbol{F}}_D^w = \hat{\boldsymbol{V}}^w, \quad [10]$$

where

$$\boldsymbol{q}^w = \left(\hat{\boldsymbol{V}}^w \cdot \hat{\boldsymbol{n}}^w\right)\left(\left(\hat{\boldsymbol{V}}^w \times \hat{\boldsymbol{n}}^w\right) \times \hat{\boldsymbol{V}}^w\right) \quad [11]$$

and $\hat{\boldsymbol{n}}^w$ is the unit vector normal to the plane of the strip in its dorsal direction (see Fig. S1E). It is imperative to note that $C_L$, $C_D$, $V$, $c$, $\hat{\boldsymbol{F}}_L$, $\hat{\boldsymbol{F}}_D$ and $\alpha_r$ are functions of $r$. Their values vary for different blade element strips along the span of the wing. Moreover, our calculation of the unit vector $\hat{\boldsymbol{F}}_L$ was sufficient to keep track of the direction of the lift force vector, without invoking a sign from the lift coefficient $C_L$ outside the range of the angle of attack from 0 to $\pi/2$ radians. In the wing-attached coordinate frame, the total translational aerodynamic force on a strip is

$$d\boldsymbol{F}_{tra}^w = d\boldsymbol{F}_L^w + d\boldsymbol{F}_D^w. \quad [12]$$

***I.4. Rotational Aerodynamic Force.*** We also calculate the aerodynamic force due to its rotation with angular velocity $\dot{\alpha}$ about the $y^w$-axis (18). This force was assumed to be acting perpendicular to a blade element strip at a distance half-chord behind the leading edge (10). In the wing-attached coordinate frame, the rotational aerodynamic force on a small wing strip of width $dr$ is

$$d\boldsymbol{F}_{rot}^w = \rho C_R V c^2 \dot{\alpha}_h \; dr \; \hat{\boldsymbol{F}}_{rot}^w \; , \quad [13]$$

$$\hat{\boldsymbol{F}}_{rot}^w = \begin{bmatrix} -\sin\alpha \\ 0 \\ -\cos\alpha \end{bmatrix}, \quad [14]$$

where the rotational aerodynamic coefficient $C_R = \pi\left(0.75 - \frac{e}{c}\right)$ and $e$ is the distance between the leading edge and wing pitching axis. $\alpha_h$ is the wing's inclination angle relative to the absolute horizontal, $\alpha_h = \alpha - \beta$, and its derivative represents the angular velocity of the wing pitching rotation with respect to the global frame (12, 17).

***I.5. Force Due to Added-Mass.*** While the wing undergoes translational and rotational accelerations during flapping, it experiences an inertial force to accelerate the boundary layer of air around the wing surface. Assuming the moth is flying at a constant velocity (on average), the most significant contributions to this force come from the wing accelerations $\ddot{\phi}$ and $\ddot{\alpha}$, and the velocity product $\dot{\phi}\dot{\alpha}$ due to the force being measured in a non-inertial reference frame. This force acts perpendicular to a blade element strip at the half-chord because the boundary layer is assumed to be uniformly distributed around the blade element strip (14). In the wing-attached coordinate frame, the force due to added-mass on a small wing strip of width $dr$ is given by the following equation (19).

$$d\boldsymbol{F}_{adm}^w = \frac{1}{4}\pi\rho\left(\left(\ddot{\phi}\sin\alpha + \dot{\phi}\dot{\alpha}_h\cos\alpha\right) r c^2 + \frac{1}{4}\ddot{\alpha}_h c^3\right) dr \; \hat{\boldsymbol{F}}_{adm}^w \; , \quad [15]$$

$$\hat{\boldsymbol{F}}_{adm}^w = \begin{bmatrix} \sin\alpha \\ 0 \\ \cos\alpha \end{bmatrix}, \quad [16]$$

where $r$ is the distance of the wing strip from the wing hinge along the wing pitching axis.

***I.6. Sum of Force Components.*** For each force (translational, rotational and added-mass), we numerically integrated the force of each strip along the wing span to determine the whole wing force. The three forces were then summed in the right wing-attached coordinate frame,

$$\boldsymbol{F}^{\mathrm{w}}_{\mathrm{right}} = \boldsymbol{F}^{\mathrm{w}}_{\mathrm{tra}} + \boldsymbol{F}^{\mathrm{w}}_{\mathrm{rot}} + \boldsymbol{F}^{\mathrm{w}}_{\mathrm{adm}}. \quad [17]$$

***I.7. Transformation to the Body-attached Frame.*** Coordinate transformations were performed by the following standard rotation matrices which represent the rotations about $x$, $y$ and $z$ axes by an angle $\xi$.

$$\mathbf{R}_x(\xi) = \begin{bmatrix} 1 & 0 & 0 \\ 0 & \cos\xi & -\sin\xi \\ 0 & \sin\xi & \cos\xi \end{bmatrix}, \quad \mathbf{R}_y(\xi) = \begin{bmatrix} \cos\xi & 0 & \sin\xi \\ 0 & 1 & 0 \\ -\sin\xi & 0 & \cos\xi \end{bmatrix}, \quad \mathbf{R}_z(\xi) = \begin{bmatrix} \cos\xi & -\sin\xi & 0 \\ \sin\xi & \cos\xi & 0 \\ 0 & 0 & 1 \end{bmatrix} \quad [18]$$

To determine how the aerodynamic forces act on the moth body, we transformed the force vector from the wing-attached frame to the body-attached frame. This was done in two steps. First, we transformed the instantaneous force vector from the wing-attached frame to the stroke-plane frame (through the wing kinematic angles $\phi$ and $\theta$) as follows

$$\boldsymbol{F}^{\mathrm{w}}_{\mathrm{right}} = \mathbf{R}_z(\phi)\mathbf{R}_x(\theta)\boldsymbol{F}^{\mathrm{w}}_{\mathrm{right}} \ . \quad [19]$$

Second, the instantaneous force vector was transformed from the stroke-plane frame to the body-attached frame (through the stroke-plane angle $\beta$, given that there is no body roll rotation) as follows

$$\boldsymbol{F}^{\mathrm{b}}_{\mathrm{right}} = \mathbf{R}_x(\beta_{\mathrm{r}})\mathbf{R}_y(-\beta)\boldsymbol{F}^{\mathrm{w}}_{\mathrm{right}}. \quad [20]$$

The overall transformation from the wing-attached frame to the body-attached frame can also be represented as a single transformation matrix $\mathbf{R}^{\mathrm{b}}_{\mathrm{w}}$,

$$\boldsymbol{F}^{\mathrm{b}}_{\mathrm{right}} = \mathbf{R}^{\mathrm{b}}_{\mathrm{w}}\boldsymbol{F}^{\mathrm{w}}_{\mathrm{right}} \ , \quad [21]$$

where

$$\mathbf{R}^{\mathrm{b}}_{\mathrm{w}} = \mathbf{R}_x(\beta_{\mathrm{r}})\mathbf{R}_y(-\beta)\mathbf{R}_z(\phi)\mathbf{R}_x(\theta). \quad [22]$$

However, for the left wing, the overall transformation is

$$\mathbf{R}^{\mathrm{b}}_{\mathrm{w}} = \mathbf{R}_z(\pi)\mathbf{R}_x(\beta_{\mathrm{r}})\mathbf{R}_y(-\beta)\mathbf{R}_z(\phi)\mathbf{R}_x(\theta). \quad [23]$$

Now total force due to both wings is

$$\boldsymbol{F}^{\mathrm{b}}_{\mathrm{total}} = \boldsymbol{F}^{\mathrm{b}}_{\mathrm{right}} + \boldsymbol{F}^{\mathrm{b}}_{\mathrm{left}}. \quad [24]$$

***I.8. Calculating Moments.*** Moments for translational, and rotational and added-mass forces were calculated separately because they have different moment arms. The translational force is assumed to act at the quarter chord while rotational and added-mass forces are assumed to act at the half-chord. Hence, moments due to the translational, rotational and added mass force on a small wing strip of width $dr$ in the body-attached frame are

$$d\boldsymbol{M}^{\mathrm{b}}_{\mathrm{tra}} = \boldsymbol{l}^{\mathrm{b}}_4 \times d\boldsymbol{F}^{\mathrm{b}}_{\mathrm{tra}} = \left(\boldsymbol{l}^{\mathrm{b}}_1 + r\hat{\boldsymbol{y}}_{\mathrm{w}} + d\hat{\boldsymbol{b}}^{\mathrm{b}}\right) \times d\boldsymbol{F}^{\mathrm{b}}_{\mathrm{tra}} \quad [25]$$

$$d\boldsymbol{M}^{\mathrm{b}}_{\mathrm{rot}} = \boldsymbol{l}^{\mathrm{b}}_2 \times d\boldsymbol{F}^{\mathrm{b}}_{\mathrm{rot}} = \left(\boldsymbol{l}^{\mathrm{b}}_1 + r\hat{\boldsymbol{y}}_{\mathrm{w}} + h\hat{\boldsymbol{b}}^{\mathrm{b}}\right) \times d\boldsymbol{F}^{\mathrm{b}}_{\mathrm{rot}} \quad [26]$$

$$d\boldsymbol{M}^{\mathrm{b}}_{\mathrm{adm}} = \boldsymbol{l}^{\mathrm{b}}_2 \times d\boldsymbol{F}^{\mathrm{b}}_{\mathrm{adm}} = \left(\boldsymbol{l}^{\mathrm{b}}_1 + r\hat{\boldsymbol{y}}_{\mathrm{w}} + h\hat{\boldsymbol{b}}^{\mathrm{b}}\right) \times d\boldsymbol{F}^{\mathrm{b}}_{\mathrm{adm}} \quad [27]$$

where $\boldsymbol{l}_4$ is a time-varying vector from the body center of mass to quarter chord of the wing strip, $\boldsymbol{l}_2$ is a time-varying vector from the body center of mass to half chord of the wing strip, $\boldsymbol{l}_1$ is the vector from the center of mass of the body to the wing hinge,

$$\boldsymbol{l}^{\mathrm{b}}_1 = l_1 \begin{bmatrix} \cos\chi_{\mathrm{wh}} \\ 0 \\ -\sin\chi_{\mathrm{wh}} \end{bmatrix},$$

the angle $\chi_{\mathrm{wh}}$ is the elevation angle of the wing hinge from the center of mass with respect to the horizontal plane as shown in Fig. S1F, $r$ is the distance of the small strip along the $\hat{\boldsymbol{y}}^{\mathrm{w}}$-axis, $d$ and $h$ are the signed position of the quarter chord line and the half chord line, respectively, with respect to the $\hat{\boldsymbol{y}}^{\mathrm{w}}$-axis. These positions are positive in the direction of the leading edge. These moments can be summed over the length of the wing to calculate the total moments $\boldsymbol{M}^{\mathrm{b}}_{\mathrm{tra}}$, $\boldsymbol{M}^{\mathrm{b}}_{\mathrm{rot}}$ and $\boldsymbol{M}^{\mathrm{b}}_{\mathrm{adm}}$. Then the aerodynamic moment on the body due to the right wing can be calculated as

$$\boldsymbol{M}^{\mathrm{b}}_{\mathrm{right}} = \boldsymbol{M}^{\mathrm{b}}_{\mathrm{tra}} + \boldsymbol{M}^{\mathrm{b}}_{\mathrm{rot}} + \boldsymbol{M}^{\mathrm{b}}_{\mathrm{adm}}. \quad [28]$$

Similarly calculating for the left wing, the total aerodynamic moment due to both wings is

$$\boldsymbol{M}^{\mathrm{b}}_{\mathrm{total}} = \boldsymbol{M}^{\mathrm{b}}_{\mathrm{right}} + \boldsymbol{M}^{\mathrm{b}}_{\mathrm{left}}. \quad [29]$$

**J. Aerodynamic Power.** Based on (20), we assumed that the aerodynamic power of flapping wings can be divided into three components: profile power, induced power and parasitic power.

***J.1. Profile power.*** The profile power is the rate of work done by a wing against the profile drag force on the wing. According to (13), for a small wing strip

$$dP_{\mathrm{pro}} = \frac{1}{2}\rho C_{\mathrm{D,pro}} V_{\mathrm{r}}^{3}\ dS, \qquad [30]$$

where

$$C_{\mathrm{D,pro}} = \frac{7}{\sqrt{Re}}, \qquad [31]$$

$$\boldsymbol{V}_{\mathrm{r}} = \boldsymbol{V} + V_{\mathrm{ind}}\hat{\mathbf{g}}, \qquad [32]$$

$\boldsymbol{V}$ is the relative airflow velocity, $\mathbf{g}$ is the vector of acceleration due to gravity, and $V_{\mathrm{ind}}$ is the induced speed. The Reynolds number was calculated as

$$Re = \frac{\rho \bar{c}\ \overline{V}}{\mu}, \qquad [33]$$

where $\rho = 1.184$ kgm$^{-3}$ is the density of air, $\bar{c}$ is wing's mean chord length, $\overline{V}$ is mean relative airflow speed, and $\mu = 1.849 \times 10^{-5}$ Pas is the dynamic viscosity of air at 25° C.

***J.2. Induced power.*** Induced power is the rate of work done by the wings to maintain enough vertical force that balances the weight of the animal, excluding the contribution to the vertical force from the profile drag. According to (13), the induced power of both wings can be estimated as

$$P_{\mathrm{ind}} = V_{\mathrm{ind}}(W - F_{\mathrm{D,pro},z}), \qquad [34]$$

where $V_{\mathrm{ind}}$ is the induced speed, $W$ is the weight of the animal and $F_{\mathrm{D,pro},z}$ is the vertical component of the profile drag force. According to (13)

$$V_{\mathrm{ind}} = \sqrt{-\frac{V_{\mathrm{b}}^{2}}{2} + \sqrt{(kV_{\mathrm{ind,0}})^{4} + \frac{V_{\mathrm{b}}^{4}}{4}}}, \qquad [35]$$

with animal's body speed $V_{\mathrm{b}}$, $k = 1.2$, and an estimate of the induced speed at hover (Rankine-Froude estimate) given in (21),

$$V_{\mathrm{ind,0}} = \sqrt{\frac{W}{2\rho\phi_{\mathrm{p\text{-}p}} R^{2} \cos\overline{\beta}}}, \qquad [36]$$

where $\phi_{\mathrm{p\text{-}p}}$ is the peak-to-peak amplitude of the stroke positional (sweep) angle, $R$ is the wing length, $\beta$ is the stroke-plane angle, and the profile drag

$$d\boldsymbol{F}_{\mathrm{D,pro}} = \frac{1}{2}\rho C_{\mathrm{D,pro}} V_{\mathrm{r}}^{2}\ dS\ \hat{\boldsymbol{V}}_{\mathrm{r}}. \qquad [37]$$

***J.3. Parasitic Power.*** We define parasitic power as the rate of work done against the drag force experienced by the body of the animal, excluding wings, assuming that all other body parts experience an equal relative airflow velocity.

$$P_{\mathrm{par}} = \boldsymbol{F}_{\mathrm{D,b}} \cdot \boldsymbol{V}_{\mathrm{b}} = F_{\mathrm{D,b}}\ V_{\mathrm{b}}. \qquad [38]$$

The body drag was calculated as

$$F_{\mathrm{D,b}} = \frac{1}{2}\rho\ C_{\mathrm{D,b}}\ S_{\mathrm{b}}\ V_{\mathrm{b}}^{2}. \qquad [39]$$

The coefficient of body's profile drag $C_{\mathrm{D,b}}$ is based on the empirical fits from data given in (13), and is a function of the angle of attack of the body, $\chi_{\mathrm{e}}$. The planform area of the body, $S_{\mathrm{b}}$, depends on body length, body width and $\chi_{\mathrm{e}}$. The data in (13) was used to estimate the expressions for both the lift and drag coefficients of the body.

$$C_{\mathrm{D,b}} = 0.977\chi_{\mathrm{e}}^{2} + 0.1, \qquad [40]$$

$$C_{\mathrm{L,b}} = 0.977\chi_{\mathrm{e}}^{2} + 1.364\chi_{\mathrm{e}}, \qquad [41]$$

where $\chi_{\mathrm{e}}$ is in radians and $-\pi/2 < \chi_{\mathrm{e}} < \pi/2$. Matlab plots of the raw data and curve fits are shown in Fig. S2.

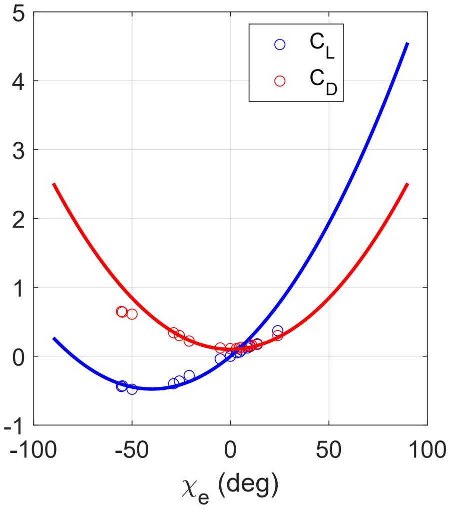


**Fig. S2.** Curve fits on the body lift and drag coefficients against the body angle of attack $\chi_e$ based on the data from (13).

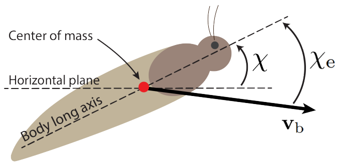


**Fig. S3.** Body angle of attack, $\chi_e$ is measured as the angle between the relative airflow on the body and the body-long axis.

Angle of attack of the body, $\chi_e$, as shown in Fig. S3, is calculated using body angle $\chi$ and the body velocity vector $\boldsymbol{V}_b$.

$$\chi_e = \chi + \tan^{-1}\frac{V_{b,z}}{V_{b,x}}. \quad [42]$$

In case $\chi_e$ is greater than $\pi/2$ or less than $-\pi/2$, it is subtracted from either $\pi$ or $-\pi$, respectively, to keep it in the range $-\pi/2 < \chi_e < \pi/2$. A moth's body is assumed to be ellipsoid with symmetrical dorsal and ventral sides.

The planform area $S_b$ of the body, assuming an ellipsoid body, is calculated as

$$S_b = \frac{\pi}{4} l_b w_b \cos\chi_e. \quad [43]$$

Because we did not have precise measurements of the body width, we estimated the body width $w_b$ from the mass $m_b$ and length $l_b$ of the body, assuming width of the ellipsoid equals depth, and the density of the body equal to water.

$$w_b = \sqrt{\frac{6m_b}{\pi l_b \rho_b}}. \quad [44]$$

**K. Trim search.** Despite choosing nearly steady wingstrokes, the kinematics of free-flying moths did not give precisely steady state aerodynamic forces from the blade-element model. Before using the aerodynamic data for analysis, we performed a trim search, i.e., we searched for the values of wing kinematic and aerodynamic parameters that created equilibrium in wingstroke-averaged forces and moments at a given flight condition. The trim search ensured three equilibrium conditions of the wingstroke-averaged forces and moment: $\bar{F}_x^b = 0$, $\bar{F}_z^b = mg$ and $\bar{M}_y^b = 0$. The remaining forces and moments $F_y$, $M_x$ and $M_z$ were already zero due to the assumption of steady forward flight in which the degree of asymmetry between left and right wings, and body motion in the lateral plane is negligible, and hence set to zero. Wing kinematic parameters included in the trim search space were $n$, $\beta_r$, and means and amplitudes of the waveforms of $\overline{\chi}$, $\overline{\beta}$, $\overline{\phi}$, $\phi_{p\text{-}p}$, $\overline{\alpha}$, $\alpha_{p\text{-}p}$, $\overline{\theta}$ and $\theta_{p\text{-}p}$. The aerodynamic parameters we included in the search were $k_D$ and $k_L$. These are scaling factors of $C_D$ and $C_L$ introduced to account for a variation in the aerodynamic lift and drag coefficients. This is reasonably expected due to slightly varying flight conditions and different wing morphology across species. The trim search for each species was performed at two forward flight conditions 1) species-averaged recorded value 2) mean forward speed of $2\,\mathrm{ms}^{-1}$ (which is roughly the mean forward speed across all wingstrokes in this study).

We set up the trim search as a computational problem that minimizes the following cost function to zero

$$G = (\bar{F}_x^b)^2 + (\bar{F}_z^b - mg)^2 + (\bar{M}_y^b)^2. \quad [45]$$

For every species, we constrained the trim search parameter space between a range of minimum and maximum values. The kinematic parameters of a species were constrained between minimum and maximum values that we measured across all recorded wingstrokes of that species in our data (Table S3 - kinematics). For all species, we constrained the parameter $k_{\mathrm{D}}$ between 0.6 and 1.4, and $k_{\mathrm{L}}$ between 0.5 and 2. These ranges were roughly based on the observation of measured variation in mean lift and drag coefficients in (13) for moth flight conditions comparable to our data. Parameters were constrained to ensure that the trimmed aerodynamics still correspond to the respective natural kinematics of every species. To solve for local minima at zero of the cost functon $G$, we used an open source MATLAB function `fminsearchbnd()` which takes the cost function, initial condition, and bounding region in the search space as inputs, and returns the values of the function and coordinates of the search space at a local minimum. However, the search is likely to end up in a non-zero local mimimum. Thus to take care of this issue, different trials were run each time initializing at a different point in the search space until 10 zero solutions were found. Then the best solution among the 10 was selected based on how close it was to the mean wing kinematics of that species and to the $k_{\mathrm{D}}$ and $k_{\mathrm{L}}$ being equal to 1. These trim conditions of the parameters are given in Table S4. Aerodynamics of all the moths trimmed within 1% of the body weight (or 1% of the body weight times radius of second moment of area, in case of pitching moment) except for *Actias luna*. In *Actias luna* the forces trimmed to within 1% but the mean pitching moment of could only trim to 28%.

### References


1. Aiello BR, et al. (2021) Adaptive shifts underlie the divergence in wing morphology in bombycoid moths. *bioRxiv*.
2. Olsen AM, Westneat MW (2015) StereoMorph: an R package for the collection of 3D landmarks and curves using a stereo camera set-up. *Methods in Ecology and Evolution* 6(3):351–356.
3. Ellington CP (1984) The Aerodynamics of Hovering Insect Flight .2. Morphological Parameters. *Philosophical Transactions of the Royal Society of London Series B-Biological Sciences* 305(1122):17–40.
4. Matthews M, Sponberg S (2018) Hawkmoth flight in the unsteady wakes of flowers. *The Journal of Experimental Biology* 221(22):jeb179259.
5. Sponberg S, Dyhr JP, Hall RW, Daniel TL (2015) Luminance-dependent visual processing enables moth flight in low light. *Science* 348(6240):1245–1248.
6. Knorlein BJ, Baier DB, Gatesy SM, Laurence-Chasen JD, Brainerd EL (2016) Validation of XMALab software for marker-based XROMM. *Journal of Experimental Biology* 219(23):3701–3711.
7. Sane SP, Dickinson MH (2001) The control of flight force by a flapping wing: lift and drag production. *The Journal of experimental biology* 204(Pt 15):2607–26.
8. Sane SP, Dickinson MH (2002) The aerodynamic effects of wing rotation and a revised quasi-steady model of flapping flight. *Journal of Experimental Biology* 205(8):1087–1096.
9. Faruque I, Humbert JS (2010) Dipteran insect flight dynamics. Part 1 Longitudinal motion about hover. *J Theor Biol* 264(2):538–552.
10. Han JS, Kim JK, Chang JW, Han JH (2015) An improved quasi-steady aerodynamic model for insect wings that considers movement of the center of pressure. *Bioinspiration and Biomimetics* 10(4).
11. Kim JK, Han JS, Lee JS, Han JH (2015) Hovering and forward flight of the hawkmoth *Manduca sexta*: trim search and 6-DOF dynamic stability characterization. *Bioinspiration & Biomimetics* 10(5):56012.
12. Cheng B, et al. (2016) Flight mechanics and control of escape manoeuvres in hummingbirds. II. Aerodynamic force production, flight control and performance limitations. *Journal of Experimental Biology* 219(22):3532–3543.
13. Willmott AP, Ellington CP (1997) The mechanics of flight in the hawkmoth *Manduca sexta*. II. Aerodynamic consequences of kinematic and morphological variation. *Journal of Experimental Biology* 200(21):2723–2745.
14. Truong QT, et al. (2011) A modified blade element theory for estimation of forces generated by a beetle-mimicking flapping wing system. *Bioinspiration and Biomimetics* 6(3).
15. Kim JK, Han JH (2014) A multibody approach for 6-DOF flight dynamics and stability analysis of the hawkmothManduca sexta. *Bioinspiration & Biomimetics* 9(1):16011.
16. Willmott AP, Ellington CP (1997) The mechanics of flight in the hawkmoth *Manduca sexta*. I. Kinematics of hovering and forward flight. *Journal of Experimental Biology* 200(21):2705–2722.
17. Ellington CP (1984) The Aerodynamics of Hovering Insect Flight 4. Aerodynamic Mechanisms. *Philosophical Transactions of the Royal Society of London Series B-Biological Sciences* 305(1122):41–78.
18. Fung YC (1969) *An introduction to the theory of aeroelasticity*. (Dover, New York).
19. Maybury WJ, Lehmann FO (2004) The fluid dynamics of flight control by kinematic phase lag variation between two robotic insect wings. *Journal of Experimental Biology* 207(26):4707–4726.
20. Ellington CP (1984) The aerodynamics of hovering insect flight. V. A vortex theory. *Philosophical Transactions of the Royal Society of London. B, Biological Sciences* 305(1122):115–144.
21. Ellington CP (1984) The Aerodynamics of Hovering Insect Flight 6. Lift and Power Requirements. *Philosophical Transactions of the Royal Society of London Series B-Biological Sciences* 305(1122):145–181.

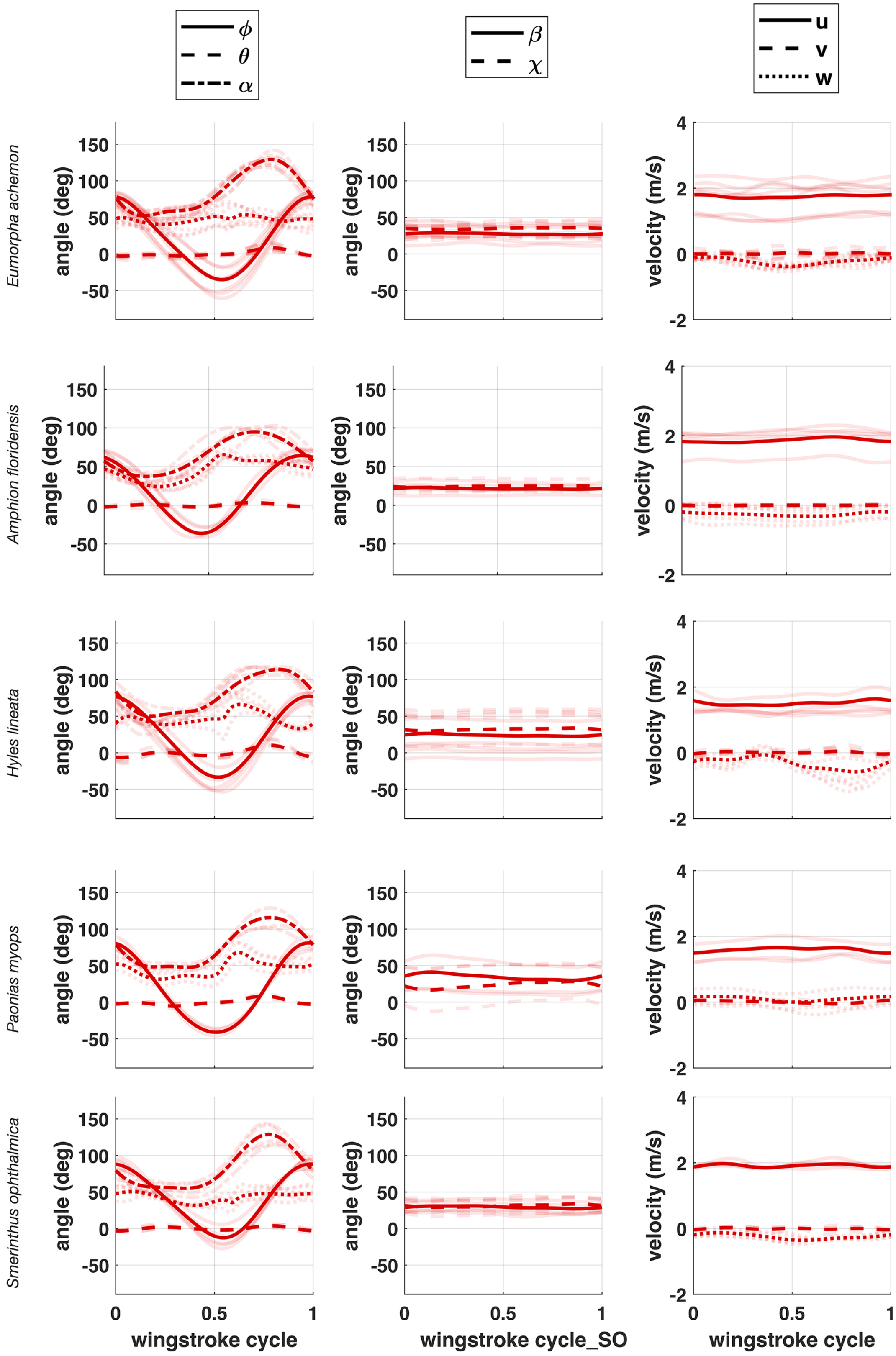


**Fig. S4.** Recorded kinematics of all hawkmoth species and inviduals.
Bold lines represent species average kinematics while transparent lines represent traces for each wing stroke across all individuals of that species. See Table S1 for symbol definitions.

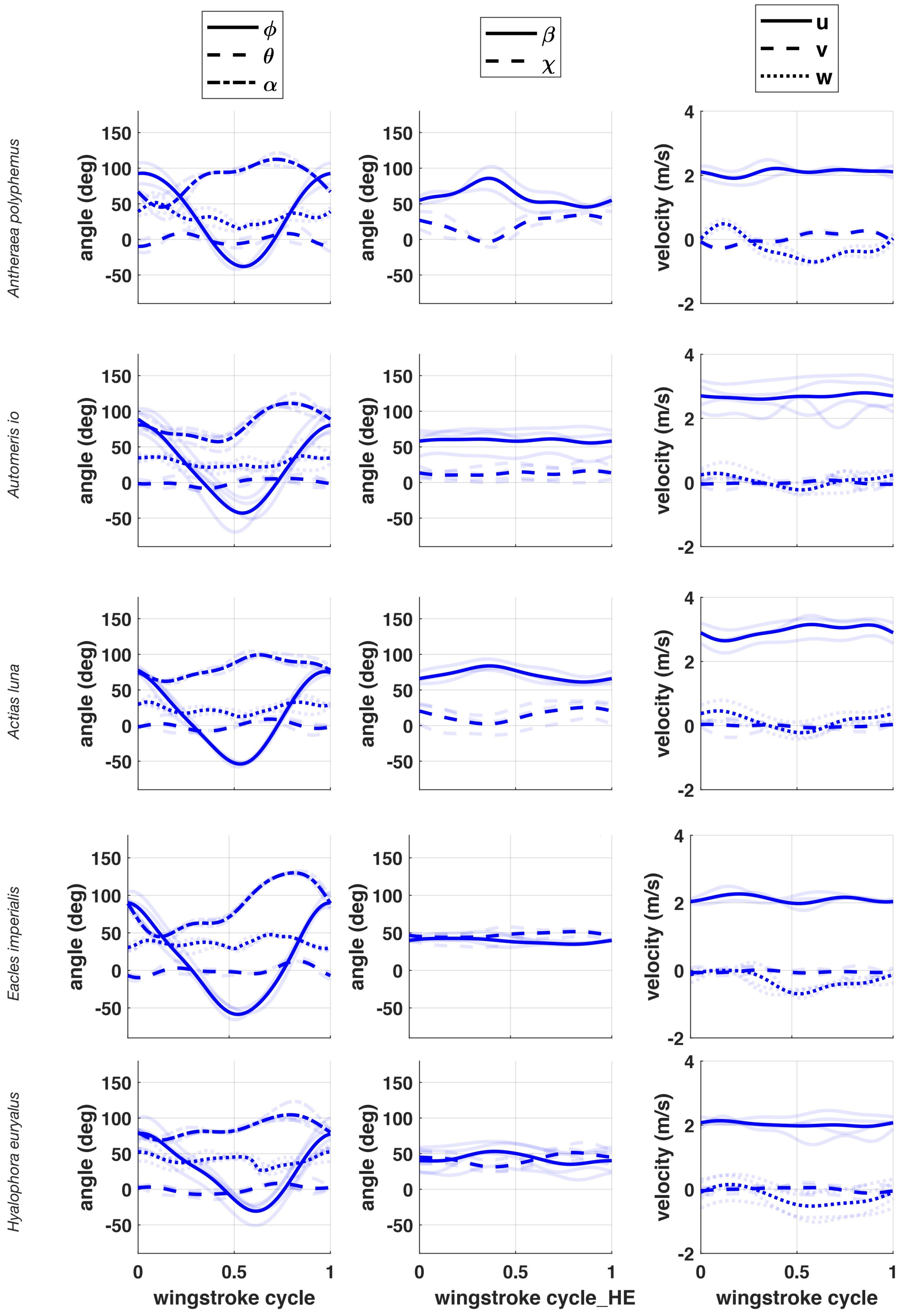


**Fig. S5.** Recorded kinematics of all silkmoth species and inviduals.
Bold lines represent species average kinematics while transparent lines represent traces for each wing stroke across all individuals of that species. See Table S1 for symbol definitions.

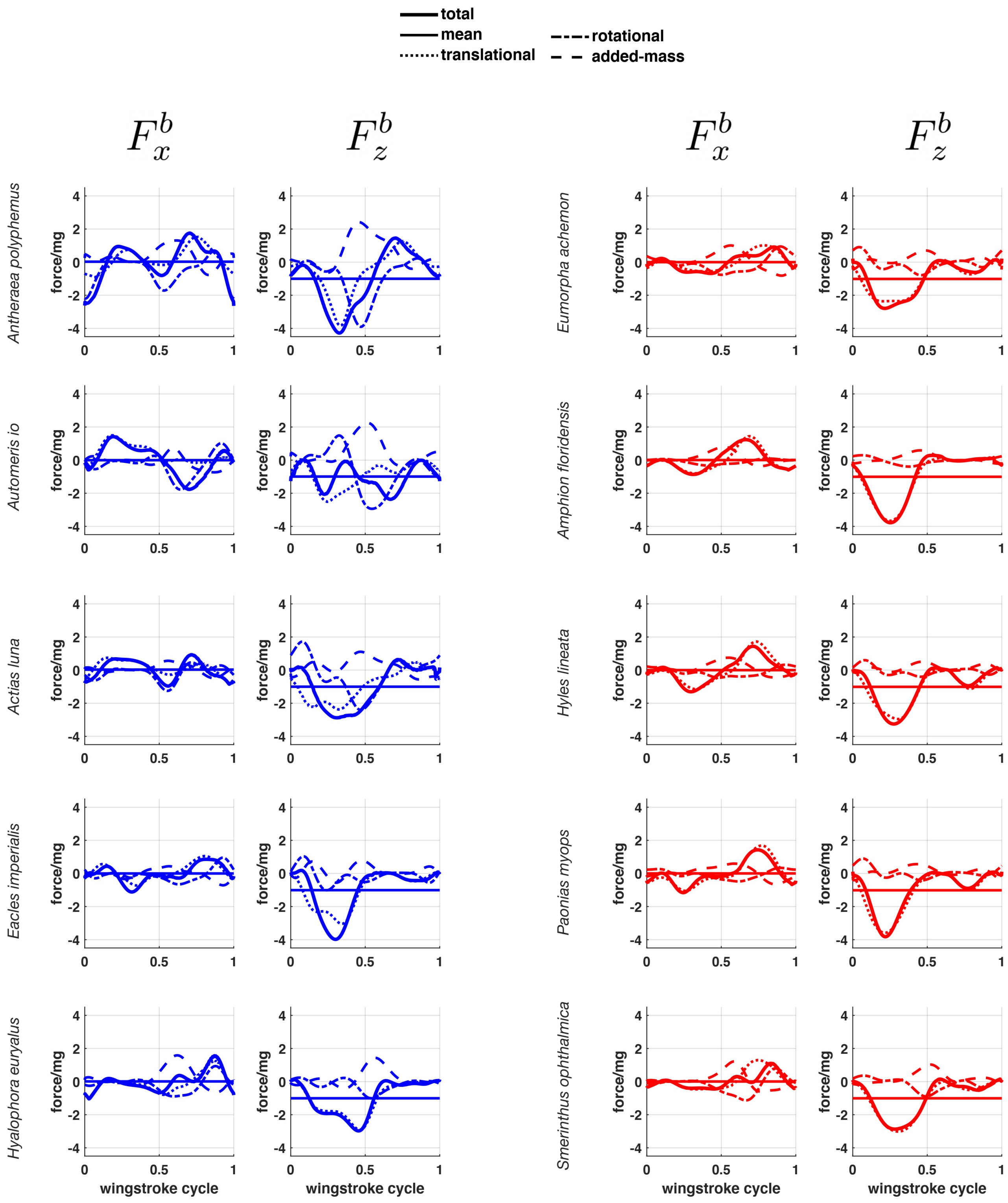


**Fig. S6.** Quasi-steady aerodynamics trimmed around each species' averaged recorded kinematics. Forces displayed are total and its components. Forces are normalized by the species mean body weight.

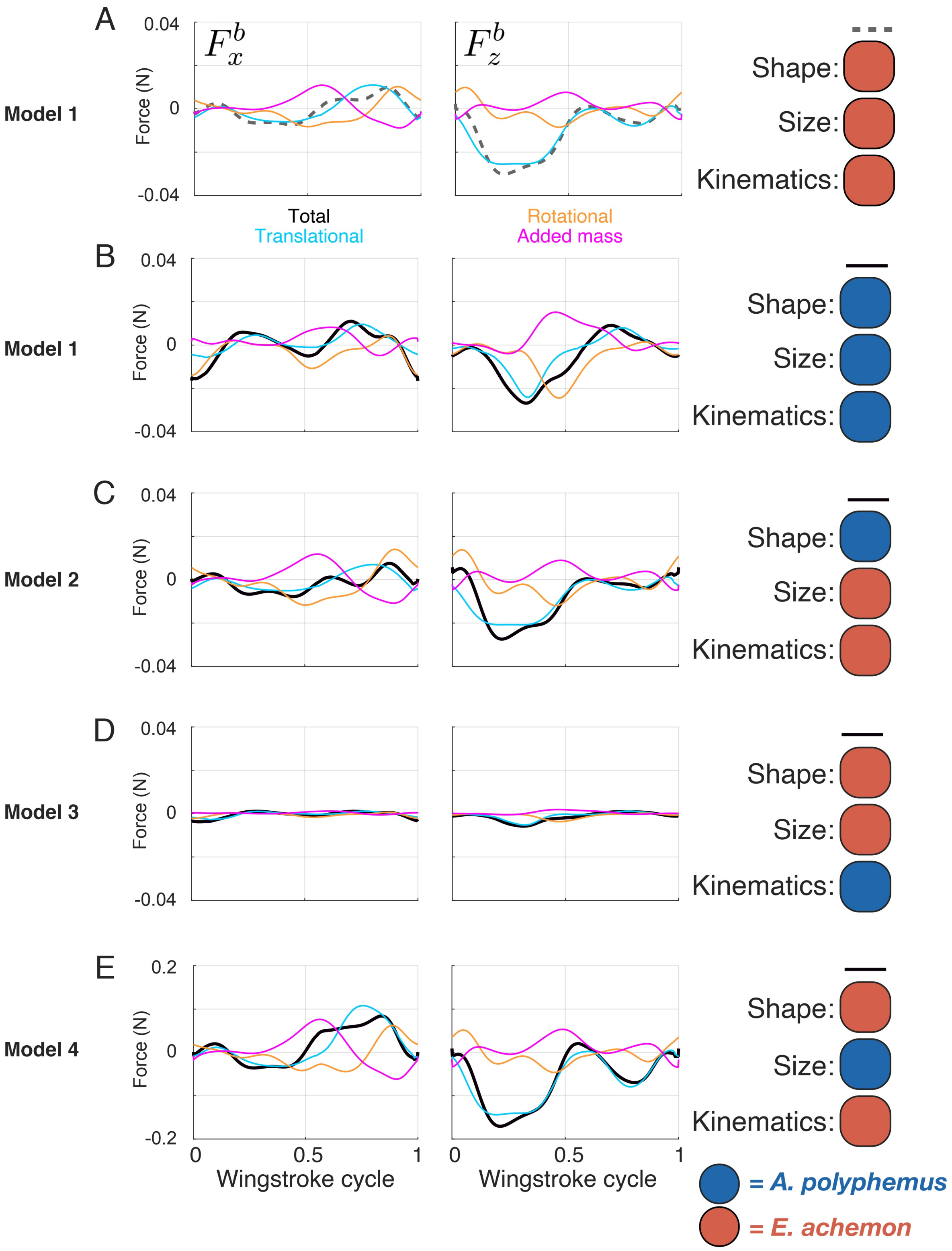


**Fig. S7.** Translational, rotational, and added mass components of aerodynamics force.
Details of the four models are identical to those in Figure 3 of the main text. Color schemes for component are the same for both species. Black represents the total force, cyan represents the translational force component ($f^b_{\text{trans}}$), gold represents the rotational force component ($f^b_{\text{rot}}$), and pink represents the added mass force component ($f^b_{\text{adm}}$). Column one and two display the $F^b_x$ and $F^b_z$, respectively. All forces are only presented for a single right wing. In all four models for each species, $F^b_{\text{trans}}$ drives the majority of the pattern in total force throughout the wing stroke.

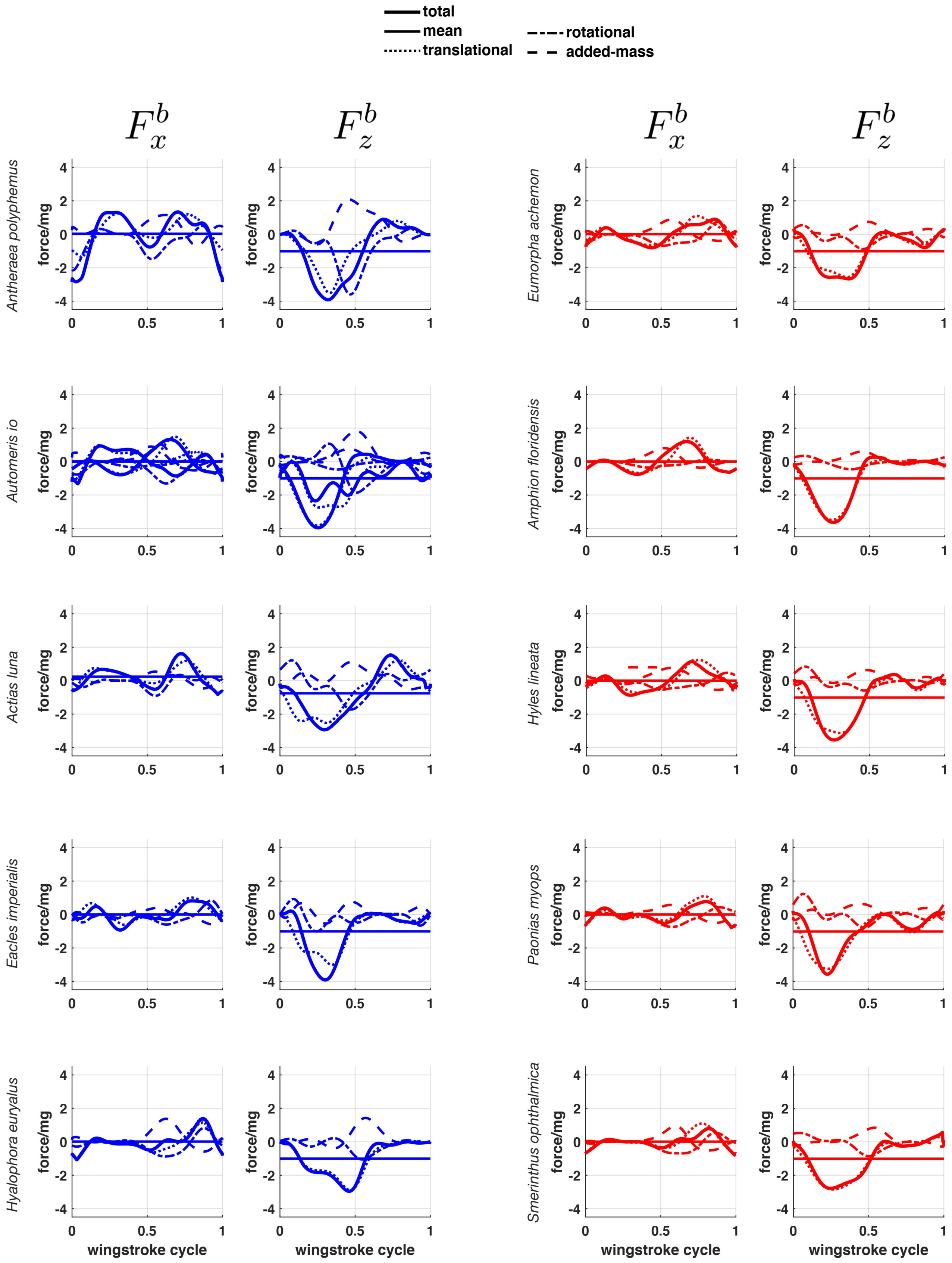


**Fig. S8.** Quasi-steady aerodynamics trimmed around each species' averaged recorded kinematics.
Forces displayed are total and its components. Forces are normalized by the species mean body weight.

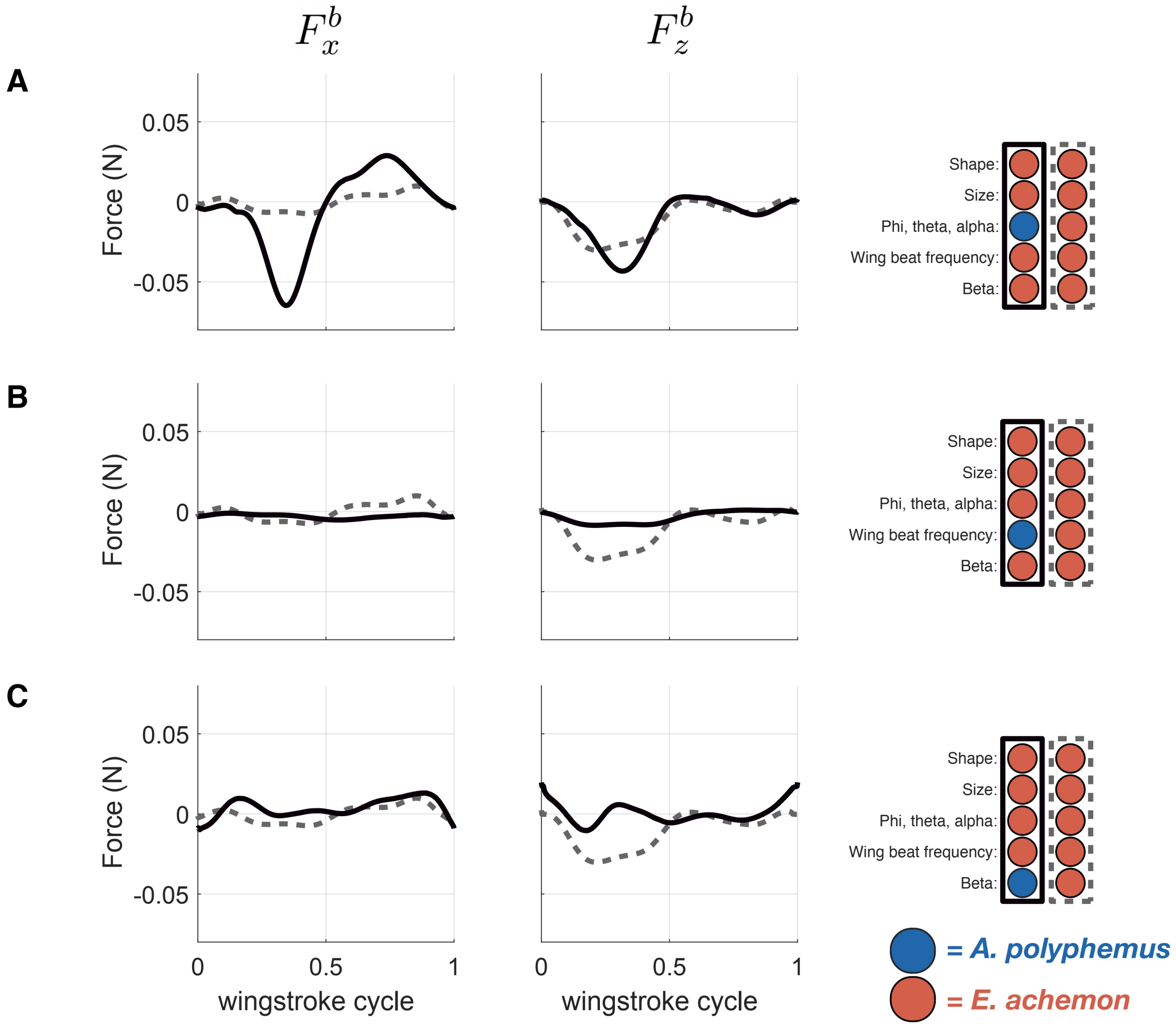


**Fig. S9.** The role of kinematic parameters in shaping the aerodynamics of each species.
This set of models investigates the contribution of wing kinematics (A), wing beat frequency (B), and stroke plane angle (C) to total aerodynamic force production. In each panel, the two models are distinguished by solid and dashed lines. The variables used in each model can be found to the right of the data and are outlined in a corresponding solid or dashed line. The color of each circle represents the species from which each variable was measured.

**Table S1. List of symbols in alphabetical order.**

| Symbol | Definition |
|---|---|
| $(a_{\phi,k}, b_{\phi,k})$ | Fourier series coefficients of Fourier fits of $\phi$ |
| $\hat{\boldsymbol{b}}$ | trailing edge to leading edge unit vector |
| $C_{\mathrm{D}}$ | aerodynamic coefficient of the drag force |
| $C_{\mathrm{L}}$ | aerodynamic coefficient of the lift force |
| $C_{\mathrm{R}}$ | coefficient of the rotational aerodynamic force |
| $c$ | chord length |
| $d$ | distance between the wing-attached $y$-axis (wing-pitching axis) and the quarter-chord line on the wing |
| $dr$ | width of an infinitesimal blade element strip |
| $e$ | distance between the leading edge and the wing-pitching axis |
| $\boldsymbol{F}_{\mathrm{total}}$ | total aerodynamic force vector |
| $\boldsymbol{F}_{\mathrm{tra}}$ | translational aerodynamic force vector |
| $\boldsymbol{F}_{\mathrm{D}}$ | drag component vector of the translational aerodynamic force |
| $\hat{\boldsymbol{F}}_{\mathrm{D}}$ | drag component unit vector of the translational aerodynamic force |
| $\boldsymbol{F}_{\mathrm{L}}$ | lift component vector of the translational aerodynamic force |
| $\hat{\boldsymbol{F}}_{\mathrm{L}}$ | lift component unit vector of the translational aerodynamic force |
| $\bar{F}$ | wingstroke-averaged force |
| $\boldsymbol{F}_{\mathrm{rot}}$ | rotational aerodynamic force vector |
| $\boldsymbol{F}_{\mathrm{adm}}$ | aerodynamic force vector due to the added mass |
| $\boldsymbol{F}_{\mathrm{right}}$ | vector of the total aerodynamic force on right wing |
| $h$ | distance between the wing-attached $y$-axis (wing-pitching axis) and the half-chord line on the wing |
| $\boldsymbol{l}_1$ | position vector from body center of mass to the wing hinge point |
| $\boldsymbol{l}_2$ | position vector from body center of mass to the half-chord line on a blade-element wing strip |
| $\boldsymbol{l}_3$ | position vector from the body center of mass to a blade element strip of the wing |
| $\boldsymbol{l}_4$ | position vector from body center of mass to the quarter-chord line on a blade-element wing strip |
| $\boldsymbol{M}_{\mathrm{tra}}$ | translational aerodynamic moment pseudovector |
| $\boldsymbol{M}_{\mathrm{rot}}$ | rotational aerodynamic moment pseudovector |
| $\boldsymbol{M}_{\mathrm{adm}}$ | aerodynamic moment pseudovector due to added-mass force |
| $\boldsymbol{M}_{\mathrm{right}}$ | total aerodynamic moment pseudovector of right wing |
| $n$ | wingbeat frequency |
| $\hat{\boldsymbol{n}}$ | unit vector normal to the dorsal surface of the wing |
| $P_{\mathrm{pro}}$ | profile power |
| $P_{\mathrm{ind}}$ | induced power |
| $P_{\mathrm{par}}$ | parasitic power |
| $P_{aer}$ | total aerodynamic power |
| $\mathbf{R}_z(\phi)$ | transformation matrix for rotation of $\phi$ radians about the $z$-axis |
| $\mathbf{R}_x(\theta)$ | transformation matrix for rotation of $\theta$ radians about the $x$-axis |
| $\mathbf{R}_y(\beta)$ | transformation matrix for rotation of $\beta$ radians about the $y$-axis |
| $\mathbf{R}_{\mathrm{w}}^{\mathrm{b}}$ | transformation matrix for rotating the coordinate system from wing-attached to body-attached frame |
| $\boldsymbol{r}$ | position vector from the wing hinge to a blade element strip of the wing along the $y^{\mathrm{w}}$ axis |
| $r$ | distance of a blade element wing strip from the wing hinge (magnitude of $\boldsymbol{r}$) |
| $t$ | time variable during a wingstroke, where $t = 0$ corresponds to the start of the downstroke |

| Continuation of Table S1 | |
|---|---|
| Symbol | Definition |
| $u$ | $x^{\mathrm{b}}$ component of the body velocity (forward speed) |
| $\boldsymbol{V}$ | relative airflow velocity |
| $\hat{\boldsymbol{V}}$ | relative airflow velocity unit vector |
| $\boldsymbol{V}_{\mathrm{b}}$ | body linear velocity |
| $\boldsymbol{V}_{\mathrm{ind}}$ | induced airflow velocity |
| $\boldsymbol{V}_{\mathrm{r}}$ | relative airflow velocity in the near field |
| $v$ | $y^{\mathrm{b}}$ component of the body velocity (side-slip speed) |
| $w$ | $z^{\mathrm{b}}$ component of the body velocity (vertical speed) |
| $x^{\mathrm{b}}y^{\mathrm{b}}z^{\mathrm{b}}$ | body-attached coordinate frame |
| $x^{l}y^{l}z^{l}$ | body-long coordinate frame |
| $x^{\mathrm{s}}y^{\mathrm{s}}z^{\mathrm{s}}$ | stroke-plane coordinate frame |
| $x^{\mathrm{w}}y^{\mathrm{w}}z^{\mathrm{w}}$ | wing-attached coordinate frame |
| $\hat{\boldsymbol{y}}^{\mathrm{w}}$ | unit vector along the wing-attached $y$-axis (wing-pitching axis) |
| $\alpha$ | wing pitching angle (feathering angle) |
| $\dot{\alpha}$ | wing pitching angular velocity |
| $\alpha_{\mathrm{e}}$ | angle of attack |
| $\alpha_{\mathrm{r}}$ | angle of attack bound between 0 and 90° |
| $\alpha_{\mathrm{h}}$ | inclination angle of wing chord relative to the absolute horizontal |
| $\beta$ | stroke-plane angle |
| $\beta_{\mathrm{r}}$ | stroke-plane roll angle |
| $\theta$ | stroke deviation angle |
| $\dot{\theta}$ | stroke deviation angular velocity |
| $\rho$ | density of air |
| $\phi_{\mathrm{p\text{-}p}}$ | peak-to-peak amplitude of the stroke positional (sweep) angle |
| $\phi$ | stroke positional angle (sweep angle) |
| $\dot{\phi}$ | stroke positional angular velocity |
| $\chi$ | body angle |
| $\chi_{\mathrm{wh}}$ | angle of elevation of the wing hinge from the center of mass with respect to the horizontal plane |
| $\chi_{\mathrm{e}}$ | angle of attack of moth's body |
| $\boldsymbol{\omega}_{\mathrm{w}}$ | wing angular velocity pseudovector due to wing kinematic motion |
| $\boldsymbol{\omega}_{\mathrm{b}}$ | body angular velocity pseudovector |
| Superscripts: | |
| b | measured with respect to the body-attached coordinate frame |
| $l$ | measured with respect to the body-long frame |
| s | measured with respect to the stroke-plane frame |
| w | measured with respect to the wing-attached coordinate frame |
| Subscripts: | |
| b | related to the moth's body |
| w | related to the moth's wing |
| $x$ | $x$-component of a vector |
| $y$ | $y$-component of a vector |
| $z$ | $z$-component of a vector |
| Accents: | |
| $\bar{X}$ | wingstroke-averaged value of $X$ |
| $\hat{\boldsymbol{X}}$ | unit vector of $\boldsymbol{X}$ |

| Species | AF | AI | AL | AP | EA | EI | HE | HL | PM | SO |
|---|---|---|---|---|---|---|---|---|---|---|
| **Force and moment values at the equilibrium** | | | | | | | | | | |
| $\bar{F}_x/mg$ | 0.001 | -0.001 | 0.019 | -0.002 | -0.001 | -0.001 | 0.000 | -0.001 | -0.002 | 0.000 |
| $\bar{F}_z/mg$ | -0.999 | -1.002 | -0.992 | -1.002 | -1.001 | -1.001 | -1.000 | -1.001 | -1.001 | -1.001 |
| $\bar{M}_y/mgr_2$ | 0.006 | 0.002 | 0.278 | 0.002 | 0.001 | 0.001 | 0.000 | 0.001 | 0.002 | 0.001 |
| **Corresponding kinematic parameter values at the equilibrium** | | | | | | | | | | |
| $n$ | 66.999 | 22.982 | 13.289 | 11.632 | 33.603 | 15.902 | 12.994 | 38.968 | 41.884 | 34.880 |
| $\bar{\chi}$ (deg) | 34.992 | 5.697 | 26.649 | 20.368 | 39.783 | 49.443 | 42.172 | 35.444 | 49.364 | 19.004 |
| $\chi_{\mathrm{p-p}}$(deg) | 2.165 | 6.203 | 23.135 | 37.706 | 2.310 | 7.451 | 20.121 | 4.156 | 11.547 | 4.555 |
| $\bar{\beta}$ (deg) | 12.000 | 66.496 | 66.000 | 67.999 | 21.459 | 35.445 | 48.066 | 13.807 | 12.446 | 38.667 |
| $\beta_{\mathrm{p-p}}$(deg) | 2.399 | 5.577 | 22.583 | 40.289 | 2.495 | 7.788 | 18.093 | 4.083 | 11.295 | 3.987 |
| $\beta_{\mathrm{r}}$(deg) | -3.459 | 0.117 | -0.459 | -4.417 | -2.620 | 0.662 | -1.827 | -0.227 | 5.642 | 2.692 |
| $\bar{\phi}$ (deg) | 25.984 | 34.901 | 14.559 | 36.328 | 12.647 | 8.648 | 36.161 | 20.533 | 18.667 | 33.016 |
| $\phi_{\mathrm{p-p}}$(deg) | 113.948 | 135.017 | 132.126 | 122.319 | 99.743 | 148.547 | 108.400 | 118.344 | 131.238 | 121.989 |
| $\bar{\alpha}$ (deg) | 67.967 | 87.991 | 84.319 | 88.213 | 80.223 | 87.449 | 91.221 | 81.581 | 77.788 | 88.711 |
| $\alpha_{\mathrm{p-p}}$(deg) | 53.019 | 65.127 | 43.913 | 45.858 | 78.190 | 84.477 | 50.715 | 67.961 | 65.616 | 73.378 |
| $\bar{\theta}$ (deg) | 1.000 | 0.623 | 0.891 | -0.240 | 0.358 | 0.051 | -0.461 | 0.339 | 0.920 | 0.239 |
| $\theta_{\mathrm{p-p}}$(deg) | 4.001 | 18.556 | 22.662 | 23.853 | 9.068 | 23.443 | 24.298 | 18.113 | 7.279 | 6.631 |
| $k_{\mathrm{L}}$ | 0.915 | 0.604 | 0.661 | 0.839 | 0.766 | 1.152 | 0.733 | 0.843 | 0.751 | 0.964 |
| $k_{\mathrm{D}}$ | 2.000 | 1.693 | 1.076 | 0.912 | 1.522 | 1.717 | 1.349 | 1.696 | 1.358 | 1.361 |

**Table S4. Results of the trim search performed at recorded body speeds.**
